# Segmenting biological specimens from photos to understand the evolution of UV plumage in passerine birds

**DOI:** 10.1101/2021.07.22.453339

**Authors:** Yichen He, Christopher R. Cooney, Zoë K. Varley, Lara O. Nouri, Christopher J. A. Moody, Michael D. Jardine, Steve Maddock, Gavin H. Thomas

## Abstract

Ultraviolet (UV) colouration is thought to be an important signalling mechanism in many bird species, yet broad insights regarding the prevalence of UV plumage colouration and the factors promoting its evolution are currently lacking. Here, we develop a novel image segmentation pipeline based on deep learning that considerably outperforms classical (i.e. non-deep learning) segmentation methods, and use this to extract accurate information on whole-body plumage colouration from photographs of >24,000 museum specimens covering >4,500 species of passerine birds. Our results demonstrate that UV reflectance, particularly as a component of other colours, is widespread across the passerine radiation but is strongly phylogenetically conserved. We also find clear evidence in support of the role of light environment in promoting the evolution of UV plumage colouration, and a weak trend towards higher UV plumage reflectance among bird species with ultraviolet rather than violet-sensitive visual systems. Overall, our study provides important broad-scale insight into an enigmatic component of avian colouration, as well as demonstrating that deep learning has considerable promise for allowing new data to be bought to bear on long-standing questions in ecology and evolution.

## Introduction

The diversity of animal colouration is among the most striking features of life on Earth. This diversity arises through selection pressures relating to, for example, signalling (social and sexual), camouflage and crypsis, thermoregulation, and parasite defence (Cuthill et al. 2017; Caro and Koneru 2021). The role of colouration in signalling is particularly complex because effective visual communication depends on both the strength of signal and perception of the receiver (Endler 1992). Selection is expected to strongly favour adaptations that maximise perception of the signal relative to background noise in the signalling environment (Endler 1993b). Fundamentally, visual communication therefore depends on the visual sensitivity of the receiver and on the light environment. The light environment itself is determined by the available light spectrum resulting from filtered solar irradiation. For example, woodland and forest canopy habitats are dominated by ambient light rich in blue and UV (Endler 1993a).

In birds visual signalling is a dominant mode of communication and diurnal birds in particular are highly sensitive to colour. However, not all birds perceive colour equally. Visual systems in birds can be classified as either violet sensitive (VS) or ultraviolet sensitive (UVS; Ödeen and Håstad 2013). The UVS cone affords greater sensitivity to UV wavelengths as well as enhanced ability to discriminate between colours. While absorption of UV is associated with darker pigments (Nicolaï et al. 2020), UV reflectance is thought to be an important signalling mechanism in many bird species (Hausmann et al. 2003; Stevens and Cuthill 2007). Despite its importance in signalling, surprisingly little is known about the distribution of UV reflectance among bird species and less still on how the combined effects of interspecific variation in visual system and light environment relates to the prevalence of UV in bird plumage. We predict that the prevalence of UV reflectance in bird plumage is higher in bird species that possess UVS visual systems, occur in regions with high levels of solar UV irradiance, and occupy primarily wooded or forested habitats that generate ambient light conditions favouring the use of UV signals for achieving conspicuousness.

Testing these predictions requires data on UV reflectance spanning species with variability in both visual system and light environment. Significant advances in our understanding of bird colouration have come from broad-scale studies that are limited to the human visual spectrum (i.e. excluding UV) (e.g. Dale et al. 2015), or include UV but are either phylogenetically limited or have sparse species sampling (e.g. Stoddard and Prum 2011; Maia et al. 2013; Cooney et al. 2019; Miller et al. 2019). However, capturing the variation to test our hypotheses requires applications of methods that capture UV reflectance across a phylogenetically broad and dense species sampling. Measuring or digitising specimens from natural history collections has become a critically important step in generating large scale data sets in ecology and evolution (e.g. Cooney et al. 2017; Felice and Goswami 2018; Sheard et al. 2020). However, processing of digitised data (e.g. specimen photographs) remains a significant and labour-intensive challenge. Deep learning offers significant potential in ecology and evolution to unlock vast amounts of data (Christin et al. 2019; Lürig et al. 2021). Here, we describe the analysis of a novel data set of calibrated images recording both visible and UV reflection and that allows objective measurements of colour. To address the processing challenge we test the efficacy of, and subsequently apply, deep learning algorithms to segment specimens and extract objective measurements of UV reflectance.

Segmentation allows measurements of the entire plumage (i.e. colour and pattern) for each specimen, facilitating measurement of multiple metrics relevant to our goal of testing the drivers of UV reflectance including mean, peak, and presence of UV colouration across the entire specimen. Segmentation is commonly used on biomedical images to separate focal regions such as cells, organs, and bones (Aljabar et al. 2009; Baiker et al. 2010; Meijering 2012) and is also beginning to be used more widely on digitised natural history datasets (Kumar et al. 2015; Unger et al. 2016). However, to be a truly scalable solution for thousands to potentially millions of images, segmentation methods must provide reliable output. We assess the performance of several traditional computer vision based segmentation methods [thresholding (Kohler 1981), region growing (Adams and Bischof 1994), Chan-Vese (Chan and Vese 2001), and graph cut (Boykov and Jolly 2001)] and compare them to semantic segmentation using deep neural networks, specifically the DeepLabv3+ architecture (Chen et al. 2018) which is from a semantic segmentation method family called DeepLab (Chen et al. 2017a; Chen et al. 2017b). The convolutional neural network (CNN) is the core deep neural network architecture for feature extraction from images (Krizhevsky et al. 2012; He et al. 2016), which takes images as input and extracts features using convolutional and pooling layers. Trained CNNs can make predictions for tasks such as image classification (Krizhevsky et al. 2012; Szegedy et al. 2014), pose estimation (Newell et al. 2016; Wei et al. 2016) and semantic segmentation (Long et al. 2015; Chen et al. 2017a) using extracted features.

Here, we assess the performance of deep learning segmentation in comparison to classic computer vision methods using photos of bird specimens taken at the Natural History Museum, Tring, UK. We then test different methods in order to build a pipeline that can segment specimen photos automatically and accurately. We used, evaluated, and compared classic and deep learning segmentation methods to segment specimens from the background and to remove obstructions (labels, string etc.) that obscure the specimen. We then generated estimates of UV signalling in bird plumage for 100’s of thousands of images from >4500 bird species using deep learning to (i) map the phylogenetic distribution of UV signalling and (ii) test how UV signalling relates to the visual system and light environment.

## Methods

### Specimen imaging

The images and labels used in this study were taken in the bird collections at the Natural History Museum, Tring. All images follow a standardised design as described by Cooney et al. (2019). We repeat the main protocols here for convenience. Photos were taken from three views (back, belly and side) for each specimen and each view was photographed twice, once in the human-visible and once in the ultraviolet (UV) light spectra, enabled by using a Nikon 105mm f/4.5 UV Nikkor lens and a modified Nikon D7000 DSLR camera. The camera was modified (by Advanced Camera Service, Norfolk; http://advancedcameraservices.co.uk/) to allow both human visible and ultraviolet (UV) wavelengths of light to be recorded. For each view, pairs of images (human-visible and UV) were taken in the human-visible or UV spectrum by using either a Baader UV/IR Cut filter / L filter (transmits light in the human visible range 400-680nm) or a Baader U-Venus-Filter (transmits light in the UV range 320–380 nm). Each image included one specimen and a set of five Labsphere Spectralon diffuse reflectance standards (2%, 40%, 60%, 80% and 99% reflectance arranged left to right in each image, referred to as Standard 1-5) photographed against a non-reflective black background (theatre blackout curtains) under controlled lighting conditions (two Bronocolor Pulso G 1600 J lamps with UV filters removed and powered by a Broncolor Scoro 1600S Power Pack). Specimens were placed with heads on the left and tails on the right in images where possible. Due to variation in size and shape of different species (e.g. exceptionally long neck or legs) some museum specimens are arranged in non-standard ways (e.g. fold necks to fit specimens in the camera). The same camera settings were used for all photographs (1/250 sec, f/16.0, ‘Daylight’ white balance, RAW photo format), with the exception that ISO was 100 for human visible images and 1000 for UV images. Images were saved in RAW format at a resolution of 4,948 × 3,280 pixels. The full data set used for deep learning consists of 122,610 visible-light images consisting of 40,870 specimens from 8,504 species (mean sampling of 2.47 male and 2.26 female specimens per species).

### Image segmentation with deep learning

We used DeepLabv3+ (Chen et al. 2018) to create a segmentation workflow with two steps: (i) data preparation, including expert labelling to generate training and model evaluation data sets, and image downsampling; and (ii) model training and application.

#### Data preparation

We produced data for model training and assessment by manually labelling a subset of 5,094 photos representing three views of 1,698 bird species. The sample of 1,698 bird species encompass representatives of more than 81% of bird genera and 27 bird orders, so the labelled images capture a large extent of the total variation in plumage colour, patterns, and bird body shape. Examples of expert labelling are shown in Figure 1. We used multiple polygons to capture unconnected areas (Fig. 1b) and nested polygons to label non-plumage areas inside plumage areas (e.g. eyes and feet; Fig. 1c). Our goal is that segmentation should not include any regions outside the plumage area, and it is preferable to segment within the focal area (i.e. to be conservative in the estimation of the plumage area) to ensure that the colour space only contains plumage colour information. The resulting manual segmentation then contains two classes: plumage areas and non-plumage areas.

**Figure 1.**
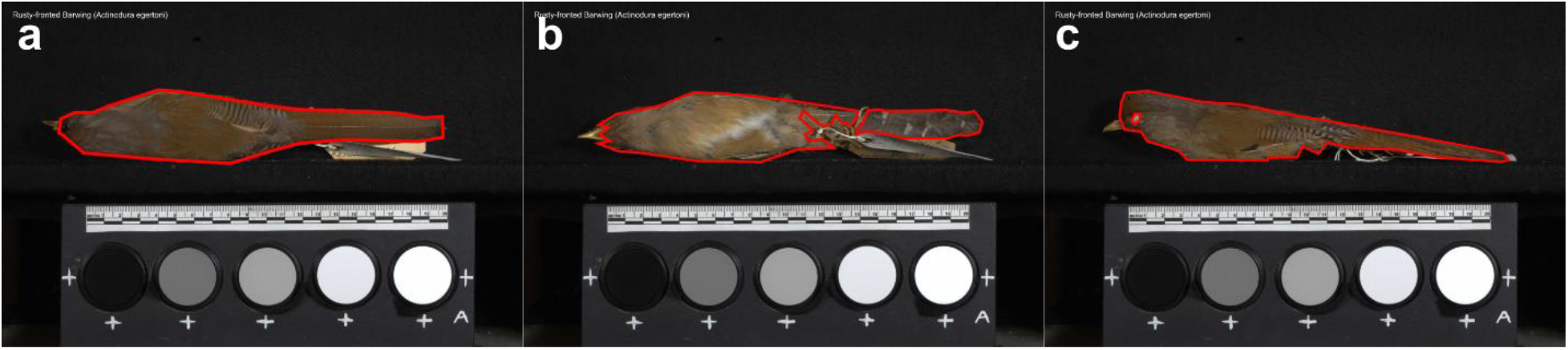
Examples of using polygons to segment plumage areas of specimens. (a) A specimen is segmented using a single polygon. (b) A specimen is segmented using multiple polygons. (c) A specimen is segmented using nested polygons as the eye is not plumage area and is excluded using a nested polygon.

DeepLabv3+ outputs heatmap arrays in which the array resolution is the same as the input image and where the number of matrices in the array is equal to the number of pixel classes. Here, we have two-pixel classes distinguishing pixels that are either inside or outside the segmented area. The output heatmap pixel value (0 to 1) of each channel represents the probability that the pixel belongs to the corresponding class. We converted coordinates of expert drawn polygons to heatmaps, with the first channel as the non-plumage area and the second channel as the plumage area. Pixels of the non-plumage area were set to 1 for the first channel and 0 for the second channel, and vice versa for pixels of the plumage area.

The DeepLabv3+ architecture is most efficient when run on a GPU but typically requires downsampling of input images to avoid memory limitations. We used a 12GB NVIDIA GTX 1080Ti GPU and downsampled all 5094 images to 618 × 410 pixels (from 4,948 × 3,280 pixels) using bilinear interpolation from the OpenCV computer vision library (Bradski 2000). This resolution is eight times smaller than the original resolution and is the largest resolution that could be trained within memory limitations.

A common approach used in many studies is to split data into a training set, a validation set, and a test set that is used to provide the final benchmark (e.g. methods in solving the ImageNet challenge; Krizhevsky et al. 2012; He et al. 2016). Here, we used only training and validation sets so that every image from the labelled dataset (covering a wide range of extant bird species) has a prediction from the same data partition routine. This allows the relationship between bird taxonomy and network performance to be evaluated (i.e. to assess whether performance varies among bird clades due to broad differences in size, shape and colouration of specimens). We split the 5094 expert labelled images into a training set and a validation set with an 80:20 ratio.

#### Model training and application

After data splitting, we trained the model with the training set (80% of images) under a set of pre-defined network hyperparameters (see below). We used five-fold cross-validation to provide an accurate estimate of model performance by averaging performance for different partitions (five partitions for five-fold cross-validation) of training and mutually exclusive validation sets. For each training step, the network generates predictions from input images. The model optimises the loss between output heatmaps and ground truth heatmaps (i.e. the expert labelled validation data set) by updating its parameters with the gradient of a loss function (Ruder 2016). We used the sum of cross-entropy between pixel values of output heatmaps and ground truth heatmaps as the loss function (Chen et al. 2017a). To minimise the loss function, we used the ADAM optimiser (Kingma and Ba 2014) and the gradient of the loss function to update model parameters. We set the initial learning rate to 0.01. Through the training process, the learning rate was cosine decayed and restarted at the initial value after reaching zero, which increases the likelihood of reaching a better local optimum (Loshchilov and Hutter 2016). The length of the first period of decay-restart was set to one epoch (defined as one pass of the full training set for the network). After each period, the new period is two times longer than the previous one (i.e. the second period takes two epochs to decay to zero, the third period takes four epochs and so on). We trained the model over 31 epochs (i.e. five complete decay-restart periods), after which the optimisation had converged (i.e. the loss has stopped decreasing).

We implemented and trained the network using Python 3 and the deep learning library Tensorflow 1.12 (Abadi et al. 2016) on one NVIDIA GTX 1080Ti GPU (12GB GPU memory). The code can be found in https://github.com/EchanHe/plumage. To balance the memory usage of the GPU and the optimisation at each step (Hinton et al. 2012) we divided training images into batches of four images. The model takes one batch per training step.

After the training process, we passed the validation images into the trained network to generate validation predictions. We then resized the predicted segmentations to the same resolution as the original images (4,948 × 3,280 pixels) and used these resized predictions, compared to the ground truth validation set to evaluate model performance (see below).

#### Additional model testing

In addition to the core pipeline above we also trained and validated the models (i) with alternative input resolution, (ii) with alternative input channels (human visible and UV), (iii) by applying image augmentation (a method that creates extra training images by manipulating the existing training images), (iv) by restricting training by image view (back, belly and side), (v) by lowering image quality, and (vi) by adjusting the size of the training set, to test if these effected the performance of the DeepLabv3+ model. The details of these tests and results can be found in the supplementary material. In the main text we focus on the core pipeline outlined above since this proved to be the best model configuration in our evaluation tests.

### Image segmentation with classic computer vision methods

To test if deep learning outperforms classic computer vision methods on this dataset, we also tested the performance of the thresholding, region growing, Chan-Vese, and graph cut methods from the OpenCV library (Bradski 2000). A weakness of some of these methods is that while they do not require any prior knowledge of the shape of the segmentation area, the region growing, Chan-Vese, and graph cut (but not thresholding) methods do require spatial information as starting values. These are usually points within the focal region. We used points within body region (2D points that are placed on specific bird body regions) as initial spatial information. We applied gaussian smoothing, a common pre-processing step to reduce noise for many classic segmentation methods (Lee 1983), prior to applying each of the four classic segmentation methods outlined below. We applied morphological close (close segmentation holes) and open (remove segmentation noises) as a global post-processing step (Haralick et al. 1987).

#### Thresholding

Thresholding segments an image by allocating each pixel to either the foreground or the background based on a pre-defined value (Kohler 1981). This value can be set either manually or automatically calculated based on image features such as the image histogram or entropy (Otsu 1979; Sezgin and Sankur 2004). For thresholding we first converted images to greyscale. Along with segmenting the plumage area, thresholding will inevitably segment parts of the reflectance standards, as standards necessarily span the majority of greyscale values. We therefore reduced the target area by selecting the most upper connected component of the image. This is possible because the specimen is always placed above the reflectance standards but requires the assumption that the segmented plumage area is not connected with other segmented parts. We tested whether using the modal pixel value of the image with a positive offset of 15 performs better than Otsu’s (1979) method and adaptive thresholding methods. We therefore used the modal pixel value to threshold images.

#### Region growing

Region growing is a method for segmenting the neighbouring pixels of an initial pixel. The classification of each neighbouring pixel depends on its similarity to the initial pixel values. Region growing methods iterate the same procedure by examining the neighbour pixels of newly segmented pixels until no more pixels can be segmented (Adams and Bischof 1994). We tried 150 ranges from different upper (even numbers from 2 to 30) and lower (even numbers from 2 to 20) boundaries for region growing. We found that the best combination is a lower boundary of 6 and an upper boundary of 30 and we use these settings for evaluation and comparison to DeepLabv3+.

#### Chan-Vese algorithm

The Chan-Vese algorithm is an active contour model designed to detect object outlines that are not defined by a gradient (Chan and Vese 2001) and is a development of the ‘snakes’ active contour models (Kass et al. 1988). The model requires a starting area within the segmentation area, which we initiated using squares of 20 × 20 pixels around points placed on the specimen and applied the algorithm for 100 iterations (Chan and Vese 2001).

#### Graph Cut

The graph cut algorithm (Boykov and Jolly 2001) treats an image as a graph where pixels are nodes. Each pixel has edges to its neighbour pixels, and edges to a source (foreground) and a sink (background) node. Weights of edges are based on pixel intensities and identities (i.e. foreground, background or to be segmented). The minimum cut cuts the graph into two subgraphs that have the largest weighted sum (Boykov and Jolly 2001). The result is the foreground subgraph defining the segmented object. For the graph cut method, we set points placed on the specimen as the foreground. The consistent setup for imaging specimens means that specimens would not be placed near the top, bottom, left and right boundaries, and would always be placed above the reflectance standards. We therefore set pixels within 20 pixels of the top, left and right edges and below the standard points as background.

### Segmentation evaluation

We used a range of metrics to evaluate the performance of both the deep learning and classic computer vision models. These focused on capturing the precision (positive predictive value) and recall (sensitivity) of the segmented areas and on assessing the reliability of colour information extracted from the segmentations. To assess the segmented areas we used the mean intersection over union (mIOU), precision, and recall metrics. The mIOU is the average IOU of all classes (e.g. plumage area and non-plumage area for the dataset). The IOU of class *i* is:

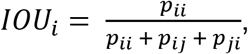

where: *p_ii_* are pixels of class *i* and classified as class *i* (true positive); *p_ij_* are pixels of class *i* but classified as other classes (false negative); and *p_ji_* are pixels of other classes classified as class *i* (false positive). IOU is a straightforward metric to measure the segmentation performance by combining aspects of both precision and recall but it can be useful to consider precision and recall separately. Precision shows the proportion of correct predictions and is a useful test of predictive capability of the model, whereas recall measures the segmentation area that the model does not predict and reflects sensitivity of the model. We used the following formulas for precision and recall of class *i*:

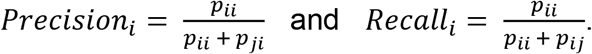

We used IOU and precision to measure the network performance as they both reflect the project-specific goal of minimising the inclusion of non-plumage regions of the image. Achieving high recall is less critical but nonetheless important because we do not want results with excessively low recall (i.e. that are extremely conservative). Segmentations have only two classes (plumage area and non-plumage area) that are mutually exclusive, so mean metrics and plumage area metrics are highly correlated. We therefore report metrics based on the evaluation of the plumage area only.

### UV data and analysis

#### Image processing

Focusing on passerine species with male and female data, all raw (.NEF) images of specimens were linearised and exported as linear TIFF files using DCRAW (Coffin 2016). Following established approaches (Troscianko and Stevens 2015; Cooney et al. 2019), pixel values were normalised using mean pixel intensity values from the five grey standards included in each image in order to control for variation in lighting conditions. We then segmented images using the image masks described above to leave only pixel values corresponding to the specimen in each image. Importantly, prior to pixel extraction each image mask was individually checked by eye and manually refined where necessary using bespoke software (https://github.com/EchanHe/PhenoLearn). The final dataset consisted of images for 24,442 specimens covering 4,545 passerine species, with an average of 2.8 male and 2.6 female specimens per species.

As individual pixel values can be noisy, and because different specimens were represented by different numbers of pixels due to their relative size in the image, we downsampled specimen images to a comparable resolution prior to extracting data on UV reflectance. To do this, we treated each specimen image as a raster and used the aggregate() function in the R package ‘raster’ (version 3.4-5) (Hijmans 2020) to find the smallest aggregation factor in the range 100 to 1 that resulted in at least 500 aggregated cells (pixels) being returned. We then randomly sampled 500 observations from this aggregated dataset to represent the plumage colouration for a particular specimen view in all further analyses.

#### Visual modelling

We used methods developed by Troscianko and Stevens (2015) to generate mapping functions to convert sampled specimen RGB pixel values into avian cone-catch values. Using tools available in the IMAGEJ Multispectral Image Calibration and Analysis Toolbox (version 2.2; http://www.empiricalimaging.com/), we generated mapping functions for each photoreceptor using equations containing second-order polynomial terms and three-way interactions between channels. Note that this approach does not incorporate information on camera responses in the UV from the camera’s green channel due to typically low sensitivities of the G channel in the UV range (Troscianko and Stevens 2015). We fit these equations to our data incorporating information on the estimated spectral sensitivities of our camera set-up and the irradiance spectrum of our illuminant (i.e. flash units), both of which we estimated previously (Cooney et al. 2019). For modelling receptor responses, we assumed idealised illumination conditions (Stoddard and Prum 2008, 2011) and receptor sensitivities corresponding to an ‘average’ ultraviolet-sensitive (UVS) avian visual system, extracted from the R package pavo (version 2.6.1) (Maia et al. 2019). We used this information to generate mapping functions for each cone class, and the resulting models were all characterised by a high degree of mapping accuracy (*R*^2^ values > 0.99). These mapping functions were used to estimate relative cone-catch values (*u*, *s*, *m*, *l*), which measure the relative contribution of ultraviolet (*u*), shortwave (*s*), mediumwave (*m*) and longwave (*l*) reflectance to plumage colour (Stoddard and Prum 2008, 2011). Our previous work has demonstrated that cone catch values generated by this photography-based approach are highly correlated (*r* > 0.92) with corresponding values calculated from spectrophotometric measurements (Cooney et al. 2019). Finally, as quantifying the colour of patches with very low overall reflectance can be problematic (Gomez and Théry 2007), pixels exhibiting a mean normalised reflectance value of <1% across all channels were re-cited to the achromatic centre (i.e. *u* = *s* = *m* = *l* = 0.25).

#### UV colouration metrics

We considered three metrics for quantifying differences in UV colouration: two based on variation in *u* values across plumages (Stoddard and Prum 2008) and a third based on determining the presence/absence of colours containing peaks of UV reflectance that may also stimulate other cone types (e.g. UV-yellow, UV-red) (Hausmann et al. 2003; Gomez and Théry 2007).

First, we calculated the average (mean) and peak (upper quartile mean) *u* values for each image. Values of *u* provide a tetrachromatic estimate of the ultraviolet contribution to plumage colouration (Stoddard and Prum 2008) and whereas mean *u* values quantify the average UV reflectance across whole plumages, peak *u* values are suitable for capturing the UV reflectance of smaller patches of colour.

Second, we employed a different approach to inferring the presence of UV colouration that involves identifying colours containing a UV peak or a peak encompassing the UV range but that may also stimulate other colour cones (Gomez and Théry 2007). This approach works by categorising colours as ‘UV colouration’ if reflectance measurements satisfy three criteria: (1) *u* cone sensitivity shows a quantum catch higher than 0.05 relative to a theoretical maximum 100% white reflectance standard, (2) reflectance over 300–400 nm exceeds 3% reflectance, and (3) reflectance over 300-400 nm is higher on average than the minimal reflectance over the range 400–700 nm (Gomez and Théry 2007). We applied these criteria to all pixel values in an image and counted the number of pixels (out of 500) subsequently categorised as a UV colour. We considered a specimen image to have evidence of UV colouration if >5% of pixels were categorised as having UV colouration.

Overall, we note that the first two metrics measure the degree to which plumage colouration exclusively stimulates the *u* cone (i.e. represents ‘pure’ UV colouration), whereas the third metric (UV colouration presence/absence) maps the occurrence of detectable peaks in UV reflectance that may occur in combination with reflectance at other wavelengths (e.g. caused by carotenoid pigmentation). We calculated estimates of each metric for each image separately, and then calculated sex-specific, species-level values for each view (i.e. body region) as the average of specimen-level values. As side view images contained large areas of plumage already captured by back and belly images (e.g. Fig. 1), we restricted our analyses to back and belly (i.e. dorsal and ventral) views only, to minimise the risk of including the same plumage area twice in our analyses.

#### Phylogenetic framework

To provide a phylogenetic framework for the passerine species included in our analysis (*n* = 4,545), we downloaded 100 trees from the posterior distribution of complete trees produced by Jetz *et al*. (Jetz et al. 2012) from http://www.birdtree.org. These trees were then pruned to generate a distribution of trees containing only the focal species set. All of our comparative analyses were run over this distribution of 100 trees to incorporate phylogenetic uncertainty into our parameter estimates. For plotting purposes, we identified a maximum clade credibility (MCC) tree from this posterior distribution of trees using the maxCladeCred() function in the R package ‘phangorn’ (version 2.5.5) (Schliep 2011).

#### Predictor variables

To test the role of factors hypothesised to influence the evolution of UV plumage colouration, we collected data for three key variables: ultraviolet-B (UVB) radiation, the degree of forest dependency, and species visual system. In total we were able to collect data on these variables for 4,527 of the 4,545 species in our dataset.

Global spatial information on annual mean UVB radiation was extracted from Beckmann *et al*. (Beckmann et al. 2014) at 15 arc-minute resolution. To generate species-level values, we intersected this dataset with information on species’ geographic ranges provided by BirdLife International (http://www.datazone.birdlife.org). To do this we first resolved taxonomic differences between the BirdLife and Jetz *et al*. datasets as far as possible, manually editing (i.e. combining or splitting) range maps for BirdLife taxa where necessary. We focused on species’ breeding geographic ranges only (seasonality = 1 or 2) and regions where species are known to be native or reintroduced (origin = 1 or 2) and extant or probably extant (presence = 1 or 2). We extracted species’ polygon range maps onto an equal area grid (Behrmann projection) at 0.5° resolution (~50 km at the equator) and then reprojected and resampled the UVB dataset to match the resolution of our range data. Species-level UVB values represent averages across their geographic range.

Forest dependency information was extracted from BirdLife International’s Data Zone (http://www.datazone.birdlife.org) and re-coded as a binary variable to facilitate effect size comparison. Specifically, species were coded as highly forest dependent (‘medium’ or ‘high’ dependency) or not (‘low’ dependency or ‘does not usually occur in forest’). In a small number of cases (*n* = 52) we filled gaps in this variable by consulting species’ records on http://www.birdsoftheworld.org.

Finally, we categorised species as having a violet-sensitive (VS) or ultraviolet-sensitive (UVS) visual system primarily using the information presented in Ödeen et al. (2011). The Ödeen et al. dataset provides approximately family-level resolution on visual system variation across passerines, and using this information we coded lineages and their constituent species as either VS or UVS based on the available data. Where species/clades were not sampled by Ödeen et al., we assumed that the visual system was the same as that of closely related lineages and/or the common ancestor. This makes sense as evolutionary switches between VS and UVS visual systems appear to be relatively rare across passerines (Ödeen et al. 2011) and birds more generally (Ödeen and Håstad 2013). However, one exception to this rule appears to be in the Maluridae (Australasian fairywrens and allies), where multiple shifts between violet and ultraviolet vision have occurred within a single genus (*Malurus*) (Ödeen et al. 2012). Therefore, for this genus we used the information in Ödeen et al. (2012) to recode species as necessary.

#### Statistical analyses

To test the relationship between UV colouration metrics and predictor variables across species, we used Bayesian phylogenetic mixed models implemented in the R package ‘MCMCglmm’ (Hadfield 2010; Hadfield and Nakagawa 2010). All models were run over a posterior distribution of 100 trees to incorporate phylogenetic uncertainty and posterior distributions of parameter estimates associated with different trees were pooled to give model estimates that incorporate phylogenetic error (Healy et al. 2014). In all cases, models were run for 110,000 iterations (sampled every 25^th^ iteration) with a 10,000 iteration burn-in. Mean *u* and peak *u* were log10-transformed prior to model fitting and all variables except UV colour presence/absence were standardised (mean = 0, standard deviation = 1) prior to model fitting to facilitate effect size comparison. For continuous response variables (mean *u* and peak *u*) we used family = “gaussian”, whereas for our binary response variable (UV colouration presence/absence) we used family = “categorical”. Correspondingly, we used two sets of standard non-informative priors: list(R=list(V=1, nu=0.002), G=list(G1=list(V=1, nu=0.002)))] for gaussian models and list(R=list(V=1, fix=1), G=list(G1=list(V=1, nu=0.002))) for categorical models. Finally, phylogenetic heritability (*H*^2^) values (Lynch 1991) were estimated by fitting intercept-only models for each variable of interest and then calculating the proportion of the total variance explained by phylogenetic effects across the posterior distribution of parameter estimates.

## Results

### Accuracy of deep learning for specimen segmentation

Across all three views, the DeepLabV3+ model achieved high IOU, precision, and recall (Fig. 2 and Table S1). The mean IOU was 93.1% (per view, back: 94.6%; belly: 91.9%; side: 92.9%), and 88.8% of the segmentations (4,525 out of 5,094) had IOU higher than 90%. The lowest IOU is 53.6%. The mean precision was 96.3% (per view, back: 96.8%; belly: 95.7%; side: 96.4%) and 97.9% of the segmentations (4,985 out of 5,094) had precision higher than 90%. The lowest precision was 70.0%. The mean recall was 96.6% (per view, back: 97.6%; belly: 95.8%; side 96.2%). No segmentation had recall lower than 50%. Less than 0.2% of the results (7 out of 5,094, per view, back: 1; belly: 4; side: 2) had recall lower than 75%, and less than 1.8% of the results (89 out of 5,094, per view, back: 13; belly: 43; side: 33) had recall lower than 90%. Four out of the worst five segmentations were caused by low recalls and all have precision higher than 85%.

**Figure 2.**
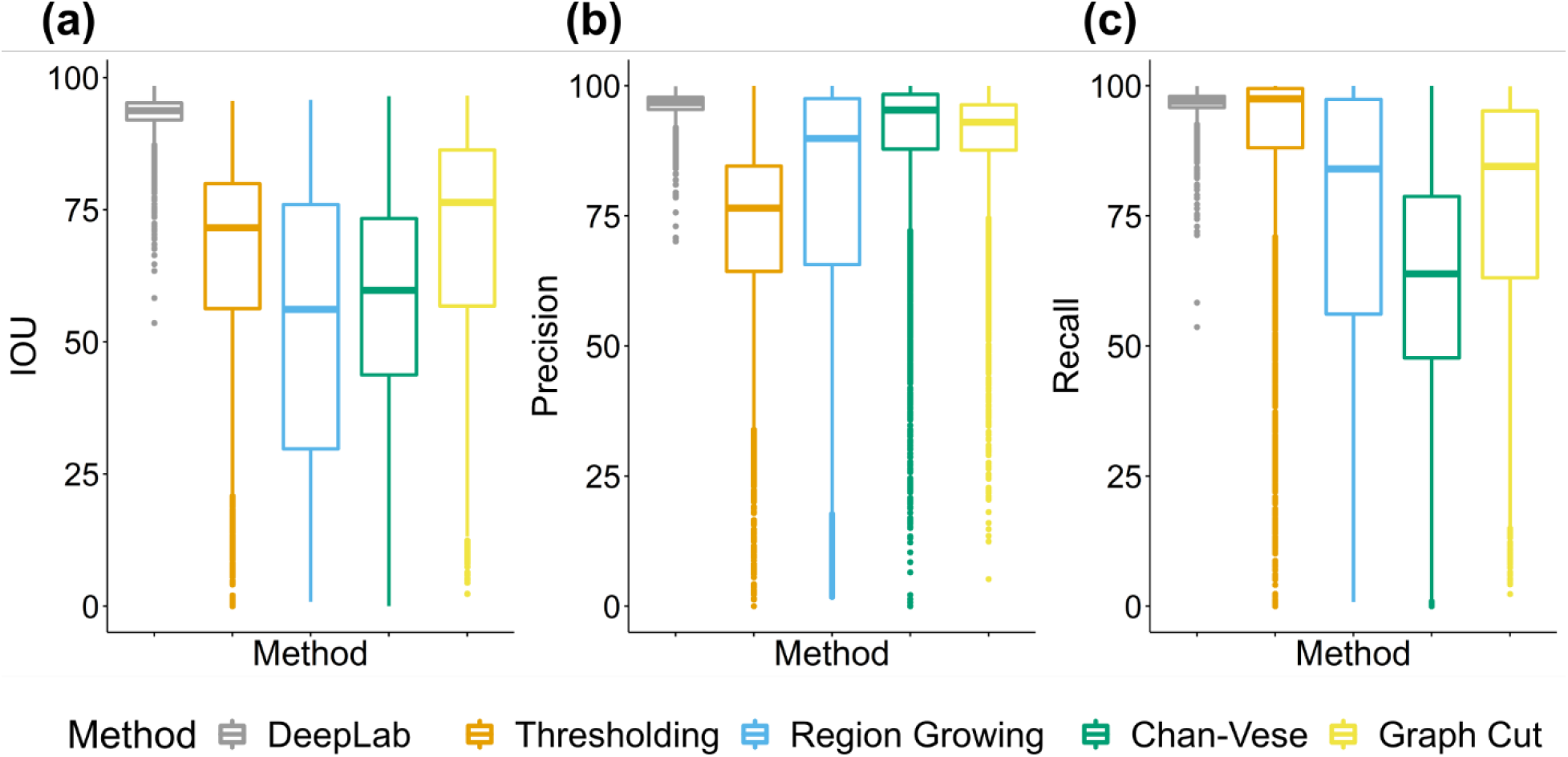
The performance of predictions (N=5,094) from DeepLabv3+ and tested classic methods (thresholding, region growing, Chan-Vese and Graph cut) for (a) IOU, (b) Precision, and (c) Recall.

Figure 3 shows the predicted deep learning segmentations on a sample of images. Many examples correctly classified eyes and labels as non-plumage area (e.g. Fig. 3a, ii-iv). Three out of four (Fig. 3a, vi-viii) of the worst IOU segmentations were caused by low recall issues (shown as large green areas in Fig. 3). The two worst recall examples (Fig. 3c, vii-viii) had many undetected plumage areas, and these images have light black backgrounds and long camera distances due to the large size of the specimens. Other low recall examples failed to detect complete tails where tails are extremely thin or irregular (Fig. 3c, v-vi). Thin tails can also cause low precision as the model misclassifies background surrounding thin tails as plumage area (Fig. 3b, vii). Legs that are placed on the top of the plumage area can be hard for the model to exclude (Fig. 3b, v). Fig. 3b, vi, also shows an example of misclassifying an irregular beak as the plumage area.

**Figure 3.**
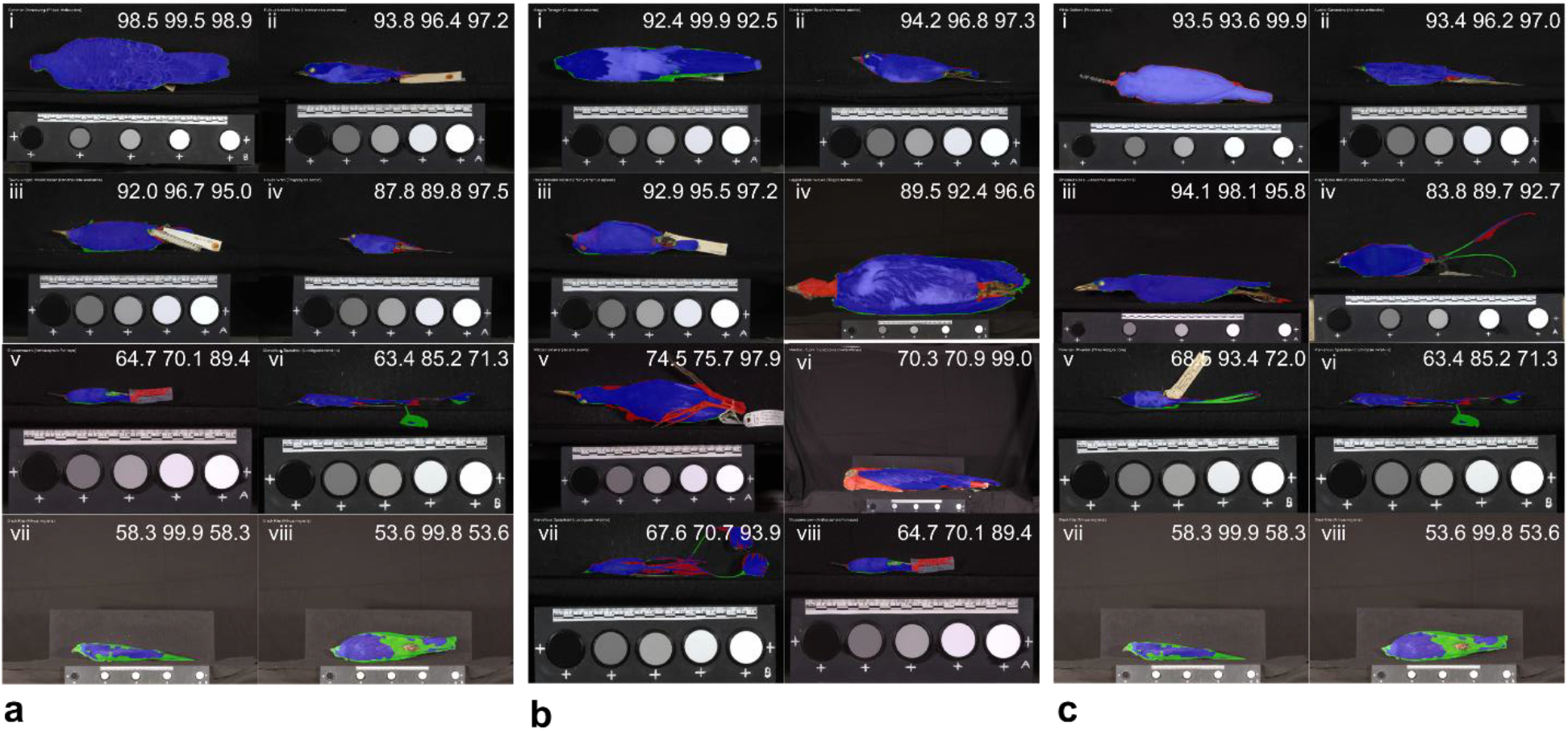
Images of the best, 50th, 75th and 95th percentile (ranked by metrics from high to low; from i to iv) and 4 worst predictions (from v to viii) based on a) IOU, b) precision and c) recall. The IOU, precision and recall (from left to right) are displayed on the top right corner of each image. Blue is correctly predicted by the model (True positive); Red is the non-plumage area that has been classified as plumage area by the model (False positive); Green is the plumage area that has been classified as non-plumage area (False Negative).

Additional model testing: We found that (i) there was a significant effect of input resolution on accuracy where low resolution can result in low accuracy (Fig. S1), (ii) the input channel using RGB has the highest performance (Fig. S2), (iii) predictions using the image augmented training set were worse than predictions using the original training set (Fig. S3), (iv) training models individually by view did not increase the accuracy (Fig. S4), (v) low-quality datasets caused slightly lower model performance but the degradations were small (Fig. S5), and (vi) the size of the training set is positively correlated with the model performance but DeepLabv3+ can achieved over 90% IOU, precision and recall using just 10% of the original training set (Fig. S6). The full details of these results can be found in the supplementary material.

### Deep learning versus computer vision for specimen segmentation

We compared the results from the DeepLabv3+ model to four classic computer vision segmentation methods (thresholding, region growing, Chan-Vese and graph cut). IOU varied significantly among segmentation methods (ANOVA: F=3141.3; d.f.=4, 25465; p<0.01), as did precision (ANOVA: F=1678.6; d.f.=4, 25465; p<0.01) and recall (ANOVA: F=1989.6; d.f.=4, 25465; p<0.01). DeepLab had superior performance for IOU, precision and recall compared to classic methods, combining both the highest mean values and lowest variance for each performance metric (Fig. 2). Specifically, DeepLab outperformed classic results by at least 23.4% on IOU, 6.4% on precision and 9.5% on recall. Graph cut had the best IOU among tested classic methods, while Chan-Vese had the best precision and Thresholding had the best recall. Graph cut was the overall best classic method in plumage images, while Chan-Vese segmented area conservatively, and thresholding tended to segment lots of non-plumage regions.

The worst examples from classic methods were clearly far worse than those from DeepLabv3+. Examples shown in Figure S7 illustrate that dark plumage, high plumage colour variability and museum specimen labels can be obstacles for classic methods whereas DeepLabv3+ segmented accurately on the same images.

### Phylogenetic distribution of UV colouration

Using manually inspected image masks produced by the DeepLabv3+ method, we mapped the phylogenetic distribution of UV colouration in passerine birds (Fig. 4). Generally, UV reflectance represents a minor proportion of avian plumage colouration, with most plumages eliciting relative ultraviolet cone-catch values (*u*) of <0.25, where cone-catch values of 0.25 would be considered the achromatic null (Stoddard and Prum 2008). Despite this general pattern, the plumages of some species are characterised by extremely high levels of ‘pure’ UV colouration, including the Purple Honeycreeper (*Cyanerpes caeruleus*) and the Hooded Mountain Tanager (*Buthraupis montana*) with peak dorsal *u* values of 0.67 and 0.62, respectively, compared to a maximal value of 0.75. Using an alternative metric (UV colouration presence/absence) that accounts for the fact that UV reflectance may co-occur with reflectance at other wavelengths (e.g. UV-red), we found more extensive evidence for UV colouration across passerines (Fig. 4), albeit with a similar pattern of phylogenetic clustering. Indeed, modal phylogenetic heritability (*H*^2^) estimates for the three UV colouration metrics we consider were all >0.80 (range 0.81 to 0.93) (Table S2), indicating that UV colouration – or a lack thereof – is phylogenetically conserved across passerines, with closely related species typically exhibiting similar levels of UV colouration.

**Figure 4.**
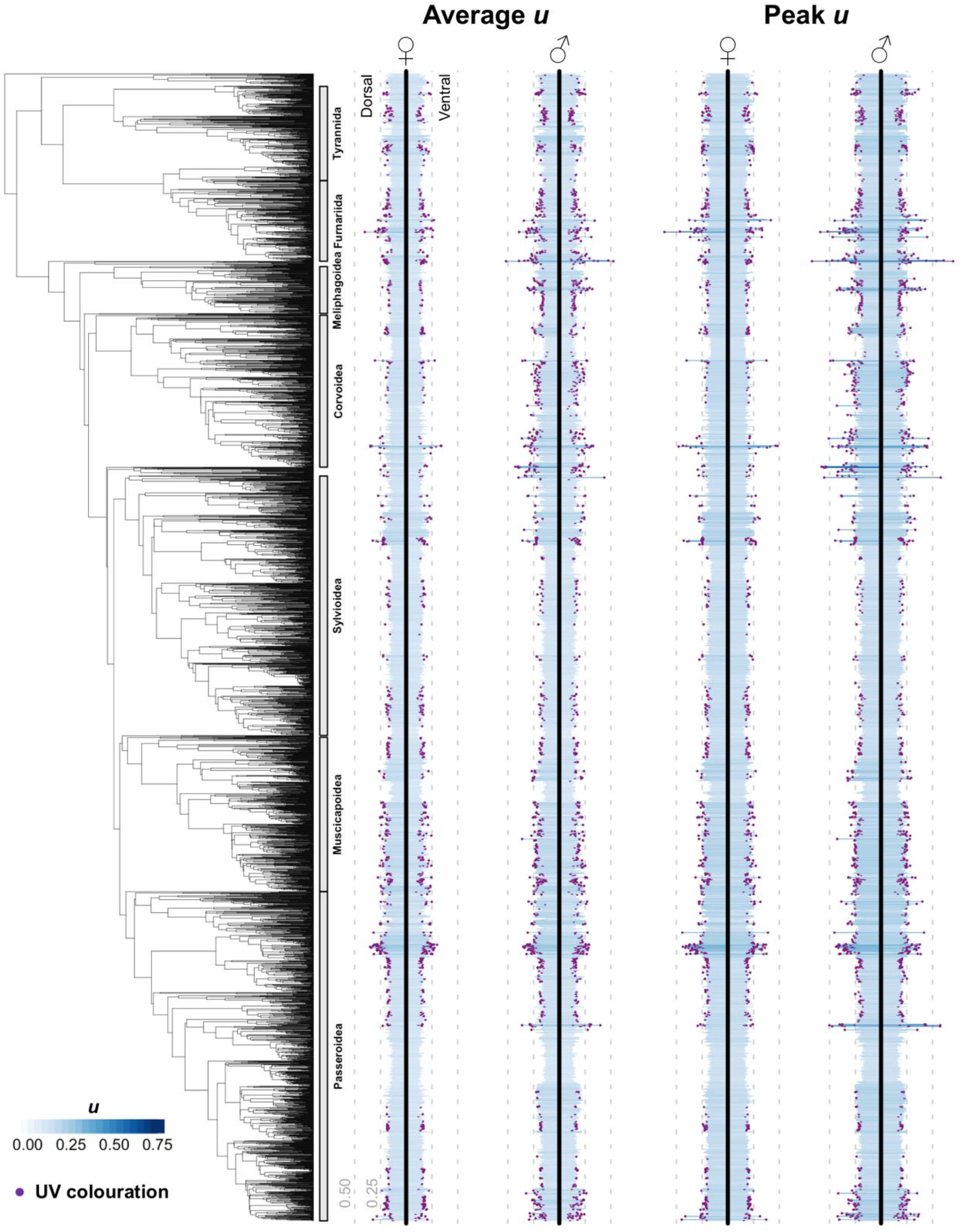
The phylogenetic distribution of UV colouration in passerine birds. Blue bars indicate the relative contribution of ultraviolet reflectance to plumage colouration (as measured by u values) of female and male individuals for 4,545 species of passerine birds. Purple dots on the end of bars (‘UV colouration’) indicate the occurrence of detectable peaks in UV reflectance possibly occurring in combination with other colours (e.g. UV-yellow).

### Correlates of UV colouration

We find that the degree of UV colouration is significantly predicted by several factors (Fig. 5, Table S3). Specifically, for average and peak *u*, we find that values are significantly higher in males, on the dorsal side of the bird, and in species inhabiting forests and locations with high incident UV radiation. Our models also revealed a notable positive association between an ultraviolet sensitive (UVS) visual system and the degree of UV colouration, but this effect was statistically non-significant and characterised by a high degree of parameter uncertainty (Fig. 5, Table S3). Results based on the UV colouration metric were similar to those based on *u* values, with the exception that UV colouration is significantly more likely to be present on the ventral, not dorsal, side of the bird.

**Figure 5.**
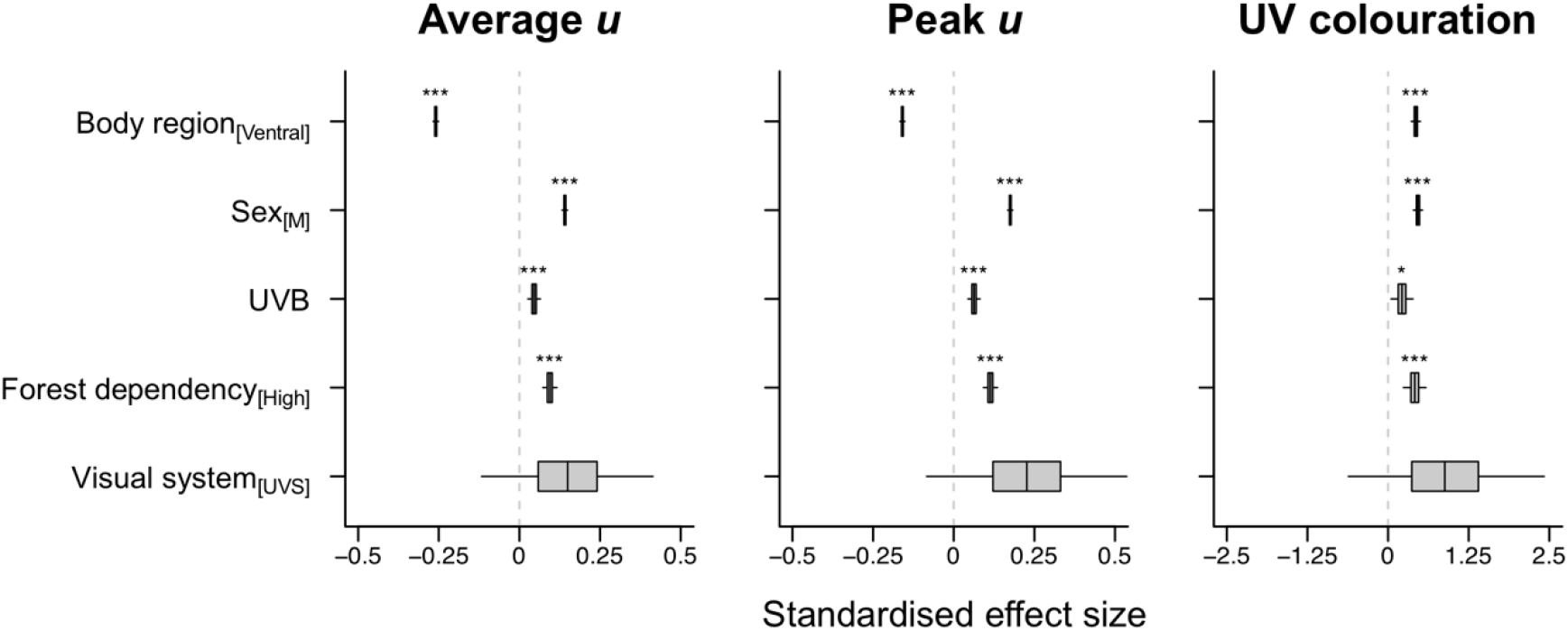
Predictors of UV colouration in passerine birds. Box plots summarise the posterior marginal distributions for all fixed-effects from Bayesian phylogenetic mixed models applied over a sample of 100 phylogenetic trees. Box widths represent the interquartile range, the median is shown as a vertical line within each box, and whiskers denote the 95% credibility interval of the distribution. Asterisks indicate evidence for a non-zero effect of the relevant variable. *, P < 0.05; **, P < 0.01; ***, P < 0.001. M, male; UVS, ultraviolet sensitive.

## Discussion

Our results show that UV reflectance, particularly as a component of other colours, is widespread across the passerine radiation. Some clades [e.g. tanagers (family: Thraupidae), corvids (family: Corvidae), thrushes (family: Turdidae)] are particularly notable for the extent of UV reflectance whereas other have comparatively low incidence or prevalence [e.g. larks (family: Alaudidae), ovenbirds and woodcreepers (family: Furnaridae)]. The presence of UV in both male and female plumage shows a strong phylogenetic signal. We also find strong evidence in support of Endler’s (1993a) hypothesis on the role of light environment on colour signal. Specifically, we find that UV (whether measured as average, peak, or prevalence as part of composite colours) is a more dominant component of plumage reflectance in species that are forest dependent and occur in regions with high levels of UVB solar radiation. UV reflectance is also notably stronger in males than females and, for mean and peak, on the dorsal, rather than ventral side of the body. These latter results in particular support the idea that UV plays an important role as a sexual signal.

Different regions of birds’ body are likely to have different roles along a crypsis to conspicuousness spectrum. Dorsal body regions have previously been shown to be more conspicuous (e.g. Delhey 2020) than ventral regions. We suggest that UV functions as a signal enhancer, increasing the conspicuousness of visual cues, and that the stronger signal in males suggests that these are cues that may often play a role in female mate choice. The signalling role of UV is implicated in the private channel hypothesis that suggests that UV can act as a special signal that is detectable by the intended receiver (i.e. conspecifics) but not others (e.g. potential predators; Hausmann et al. 2003; Stevens and Cuthill 2007). An important corollary is that the predominant predators of many passerine birds are birds of prey that are also sensitive to UV (Stevens and Cuthill 2007). Birds of prey possess VS rather than UVS receptors, and are therefore less sensitive to UV than passerines (Håstad et al. 2005) suggesting that UV signalling may still be a low cost way of enhancing signalling efficiency. Our analyses suggest a weak trend towards higher UV plumage reflectance among species with ultraviolet rather than violet-sensitive visual systems, in line with results based on visual modelling predicting only weak relationships between visual system variation and plumage colouration (Lind and Delhey 2015; Lind et al. 2017). However, we are necessarily cautious in this interpretation because there is wide uncertainty in parameter estimates for the effect of VS/UVS in our models. This uncertainty likely stems from the relatively low number of transitions between visual systems, and conservatism within clades, in the passerine radiation (Ödeen and Håstad 2013).

Taken together, our analyses reveal the diversity and extent of UV reflectance in passerine birds and provide new insight into the factors that underpin the ubiquity of UV colours in the avian colour gamut. The data on which these inferences lie rely on efficient processing of a vast quantity of raw input. We were able to achieve this using deep learning after first testing the suitability for these methods. We show how DeepLabv3+ can automatically segment bird plumage areas from other parts across more than 120,000 images within a few days on a consumer grade GPU and can identify the plumage area (precision: 96.3%) and plumage area completeness (recall: 96.6%) reliably. Our analysis showed that segmentation using DeepLabv3+ strongly outperformed all classic computer vision methods. Indeed, segmentations from classic methods are frequently so poor that they would often be unusable for downstream analyses of colour. Of the classic methods, Graph cut had the best average plumage area IOU but was 23.4% worse than the average IOU from DeepLabv3+. In contrast to the DeepLab predictions, images with dark birds and prominent label tags could not be reliably segmented using classic methods. Dark birds were normally under or over segmented, and label tags were included as plumage area (e.g. Fig. S7). Besides deficiencies shown in these examples, setting starting parameters for classic methods, for example choosing threshold values for thresholding and region growing by hand-crafted image features, is a troublesome task (Chang and Li 1994; Fan et al. 2001). We suggest that deep learning is likely to be of wider value for high throughput processing of very large image datasets and supports growing recognition of the potential value of deep learning for many applications in biodiversity science (Christin et al. 2019; Lürig et al. 2021).

Our experimental configurations also allow us to identify limitations and possible ways to further improve model performance for deep learning. We found that input image resolution had positive effects on performance, as expected and previously reported for DeepLabv2 (Chen et al. 2017a). In contrast, image augmentation, using additional channels and subsetting models did not improve the performance of the deep learning model. The best performance overall was achieved with DeepLabv3+ and an input resolution of 618 × 410 pixels. This resolution was the maximum we could achieve with available resources but could be increased with a more powerful GPU and we would expect that performance can therefore be improved further. Our results are also consistent with previous studies showing that the training set size is positively correlated to the model performance (Joulin et al. 2015; Hestness et al. 2017). However, small training set sizes did not decrease the performance drastically. It is possible to use just 15% of the original dataset (~600 images) to generate segmentations with 90% IOU on 1,018 validation images. This is still much more accurate than results using any of the classic method segmentation methods. The highly consistent imaging layout in our data may reduce the size of training data needed to get an acceptable result from deep learning.

The consistency of imaging in our data may partly explain the quality of performance of the deep learning model. The IOU in our best configuration was 93% which is higher than DeepLabv3+’s performance (mIOU: 89.0%) on the standard PASCAL VOC 2012 data set (Chen et al. 2018). In contrast to the PASCAL dataset, (i) our dataset has only two classes (plumage and non-plumage) while the PASCAL dataset has 21 classes (Everingham et al. 2015) and (ii) our images consist of few and fixed focal objects (one) under a consistent, high resolution imaging setup. In contrast, the PASCAL images are more varied (e.g. different objects, backgrounds). While there are specific challenges in removing unwanted parts of the images (including eyes and specimen labels), these do not seem to significantly impact model performance. These two factors may explain why no improvements were observed with image augmentation, additional channels and subsetting models, as the model had already been well trained using the highly standardised original dataset. Modern pipelines for museum collection digitization typically follow similarly consistent standards such as uniform specimen placements, background and light environment (Hudson et al. 2015; Unger et al. 2016; Hussein et al. 2020) suggesting that such data can be analysed with deep learning. However, high standard digitisation is time-consuming. We simulated low-quality images and they did not provide excessively inaccurate predictions, and the worst performance was much better than classic methods’ results (i.e. Fig. S5). This result, along with promising results on low consistent datasets such as PASCAL VOC 2012 (Chen et al. 2018), shows that the DeepLab model is likely to be robust on less consistent datasets.

Here, we have tested and applied deep learning approaches for semantic segmentation to reveal the prevalence and predictors of UV plumage colouration across bird species. However, deep learning has broader potential applications for image processing including species identification and key point placement (e.g. landmarking for geometric morphometrics). Some tasks may require larger training sets than we have used. For example, DeepLab (Chen et al. 2017a) used a training set size of 1400 images in PASCAL VOC 2012 and 2975 images in Cityscapes (Cordts et al. 2016). Tasks like classification and pose estimation have used even larger datasets, such as 1.2 million training images in ImageNet classification (Deng et al. 2009) and more than 28,000 images in MPII pose estimation challenge (Andriluka et al. 2014). Such large training sets can be generated through citizen science projects, such as the ‘Zen of Dragons’ (https://www.zooniverse.org/projects/willkuhn/zen-of-dragons). Regardless of the source of training data, all automated methods are likely to be imperfect and, depending on the goal of the project, may require expert error checking prior to downstream analysis as we used here. Nonetheless, we support the view that deep learning has great promise (Christin et al. 2019; Lürig et al. 2021)—particularly in the mobilisation of digitised images (both 2D and 3D) from natural history collections—allowing new data to be brought to bear on key outstanding questions in ecology and evolution.

## Acknowledgements

We thank M. Adams, H. van Grouw, R. Prys-Jones, and A. Bond from the Bird Group at the NHM, Tring for providing access to and expertise in the collection.

## Authors contributions

Y.H., S.M., C.R.C. and G.H.T designed the research; C.R.C., Z.K.V., L.O.N., C.J.A.M. and M.D.J. collected the image data; Y.H. and C.R.C. conducted the analyses. Y.H. wrote the manuscript, with input from all authors.

## Data and materials availability

All data is available in the manuscript or the supplementary materials. Analysis code is available from the corresponding author upon request.

## Supplementary material

### Additional model testing

#### (i) Effects of input resolution on the performance

DeepLab has been shown to perform better when using the original input resolution compared to resized resolutions (Chen et al. 2017a), in particular downscaled images result in lower accuracy for pose estimation and classification (Kim, Kwon Lee, and Mu Lee 2016). We compared resolutions that were 8, 10 and 16 times lower than the input images (i.e. 618 × 410, 494 × 328 pixels and 309 × 205 pixels) to test whether performance degrades at lower resolutions.

There was a significant effect of input image resolution on IOU (ANOVA: F=1361.0; d.f.=2, 15279; p<0.01), precision (ANOVA: F=1069.2; d.f.=2, 15279; p<0.01) and recall (ANOVA: F=456.3; d.f.=2, 15279; p<0.01) (Fig. S1). The IOU and recall of 618 × 410 pixels and 494 × 328 pixels were not significantly different from each other, while the rest of the accuracies (IOU, precision and recall) were positively related to the input resolution (Fig. S1). The image resolution of 618 × 410 pixels had the best overall performance, while the 309 × 205 pixels had the worst performance.

#### (ii) Effects of input channels on the performance

Previous studies have included non-visible light (e.g. UV and IR) information as the input in deep learning tasks, sometimes leading to better performance when compared to using only RGB channels (Basu et al. 2015; Potena et al. 2017; Milioto et al. 2018). Our dataset includes two sets of images, one filtered to include only human visible (RGB) wavelengths and one to include only UV wavelengths, because bird plumage frequently includes UV reflecting regions. All images were taken against a black background made of theatre blackout curtains with very low reflectance of the UV light. The specimens should therefore reflect more UV light than the background. To test whether the inclusion of UV improved network performance, models were trained with (i) images using RGB channels only, (ii) images using UV channels only and (iii) images using RGB plus UV channels.

There was a significant effect of input channels on IOU (ANOVA: F=395.6; d.f.=2, 15279; p<0.01), precision (ANOVA: F=184.9; d.f.=2, 15279; p<0.01), and recall (ANOVA: F=236.6; d.f.=2,15279; p<0.01) (Fig. S2). RGB was consistently better than UV and RGB+UV although the effects tended to be small (evaluation results of UV and UV+RGB were <2% worse than RGB) (Fig. S2).

#### (iii) Effects of image augmentation on the performance

Image augmentation is a common technique that increases the size of the training set by creating new labelled training images from manipulating the existing images and their labels, and has been shown to improve the model performance of DeepLab (Chen et al. 2017a). We created an augmented training set from the original training set in which images and their segmentations were randomly rotated (−15°to −1°, 1°to 15°), translated in both x and y axes (100 to 500 pixels), scaled (0.1 to 1.1). We used the augmented dataset to train the model with evaluation performed on the original validation set.

IOU was significantly higher (t_[10186]_=5.90, p<0.05) for the original dataset (Mean=93.1, Standard deviation (SD)=3.24) than the augmented dataset (Mean=92.7, SD=3.35) (Fig. S3). We also found a significant difference in the precision (t_[10186]_=6.63, p<0.05) and again the original dataset (Mean=96.3, SD=2.38) outperformed the augmented dataset (Mean=96.0, SD=2.56) (Fig. S3). However, there was no significant difference in the recall (t_[10186]_=1.81, p=0.07) (Fig. S3).

#### (iv) Effects of subsetting models on the performance

In our core pipeline, we used one deep learning model on images from all views (all-views model), but image variations of different views may introduce difficulties for the network to learn. We therefore tested the impact of training and validated separate models for each of the three image views (back, belly and side). This reduces the input data for each model run to 1,698 images (compared to 5,094 images).

We found that subsetting models by image view (i.e. back, belly, side) was significantly worse than using the all-views model, except for recall on the side view (t_[10186]_=1.43, p=0.15) (Fig. S4). The back view had the largest IOU difference (the all-views model has 0.7 higher IOU than individual models) and recall difference (recall of the all-views model is 0.5 higher), while the side view had the largest precision difference (precision of the all-views model is 0.4 higher) (Fig. S4).

#### (v) Quality of the training data

In our case, specimen images were taken in a highly consistent manner by controlling the placement of the specimen, light environment and background (Hudson et al. 2015). However, not all image datasets are likely to be so consistent due to practical limits (e.g. inadequate lighting). We tested whether greater variability in data quality could limit performance by generating artificial lower quality datasets. To do this, we applied a series of image manipulations in which (i) images were rotated (angles between −45° to 45°), translated (−500 to 500 pixels on x and y axes) and scaled (scale ratio from 0.8 to 1.2), (ii) 50% of images were randomly horizontally flipped, (iii) images were given new contrast and brightness (α from 0.5 to 2 and β from −50 to 50) using brightness and contrast adjustment functions in OpenCV (Bradski 2000; Bradski and Kaehler 2008), and (iv) a combination of manipulations from (i), (ii) and (iii). We applied these operations to both training images and validation images (in contrast to image augmentation outlined above where we did not manipulate the validation set).

Low-quality datasets had a significant negative effect on the IOU (ANOVA: F=205.3; d.f.=4.0, 25465; p<0.01), precision (ANOVA: F=132.8; d.f.=4.0, 25465; p<0.01) and recall (ANOVA: F=88.0; d.f.=4.0, 25465; p<0.01) (Fig. S5). The original dataset produced more accurate results than low-quality datasets (Fig. S5a). The 4th dataset (the combination of translation, rotation, scale, horizontal flip and manipulations of brightness and contrast) had the worst performance and examples of its predictions are shown in Figure S8). The 4th dataset was 1.9%, 1.2% and 0.9% worse than the original dataset on IOU, precision and recall (Fig. S5b).

#### (vi) Training dataset size

We manually labelled 5,094 images for this study. However, the number of labelled images may be limited by time and resources for other projects and studies. Here, we investigated the impact on deep learning accuracy using smaller training sets. Previous studies suggest that larger training set sizes may improve the performances of deep learning models (Joulin et al. 2015; Hestness et al. 2017). We used a subset of 1,018 images (20% of the dataset) as the only validation set for every result in this section. The training set (4,076 images) was randomly sampled five times for one proportion selected from 15 proportions (1%, every 5% from 5% to 50% and every 10% from 50% to 90%).

We found that model performance was positively related to the training set size following an approximately logarithmic pattern (Fig. S6). At least 10% of the dataset was required to attain IOU higher than 90%, at least 5% of the dataset to get precision and recall higher than 90%, and 15% of the dataset for precision and recall higher than 95%. With 100% of the dataset used for training, the model achieved 93.3% for IOU, 96.3% for precision and 96.8% for recall (Fig. S6).

**Figure S1.**
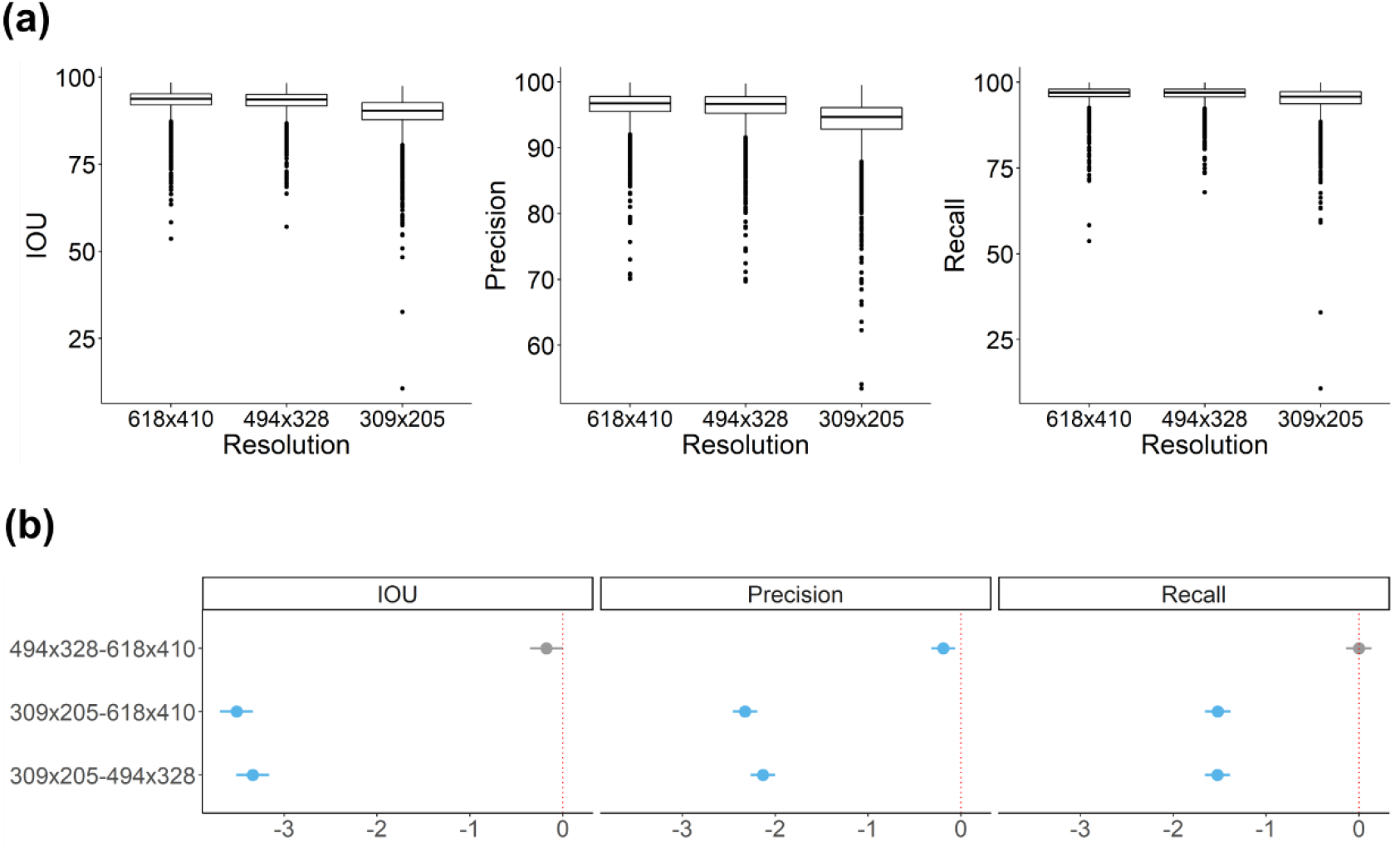
(a) Boxplots of the performances (IOU, precision and recall) of predictions (N=5,094) of tested input resolutions (618 × 410, 494 × 328 pixels and 309 × 205 pixels). (b) Plots of Tukey’s test (95% family-wise confidence level) on whether metric (IOU, precision and recall) differences among tested resolutions are significantly different (blue: significance; grey: no significance) from 0 (red dotted lines).

**Figure S2.**
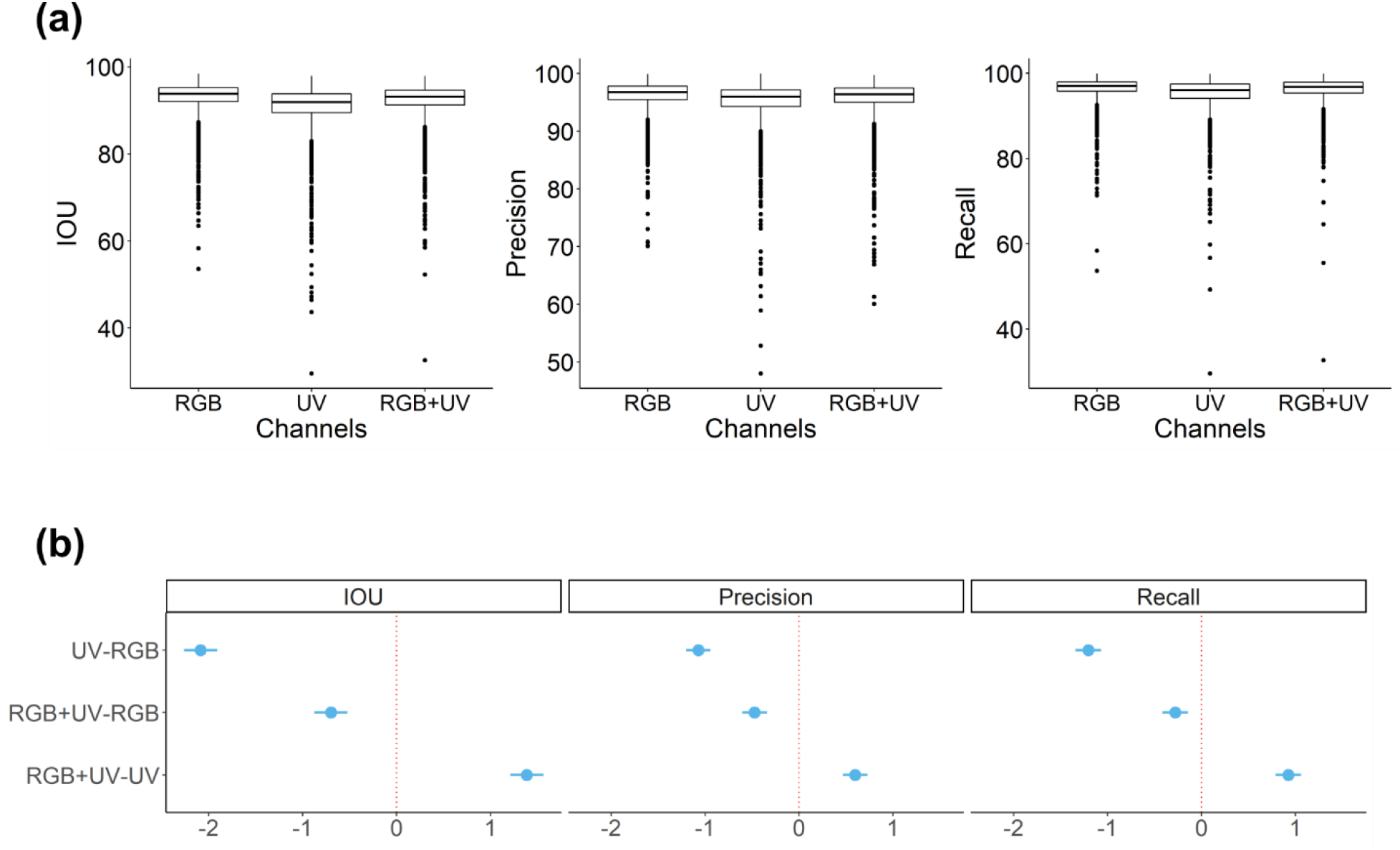
(a) Boxplots of the performances (IOU, precision and recall) of predictions (N=5,094) of tested input channels (RGB, UV and RGB+UV). (b) Plots of Tukey’s test (95% family-wise confidence level) on whether metric (IOU, precision and recall) differences among tested channels are significantly different (blue: significance; grey: no significance) from 0 (red dotted lines).

**Figure S3.**
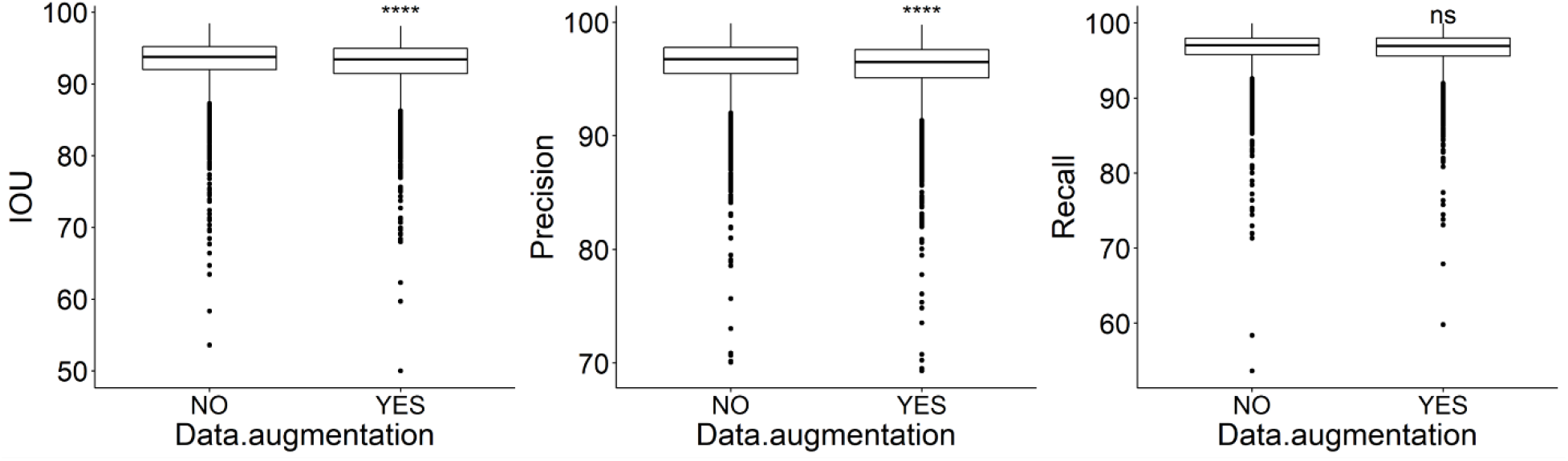
Boxplots of the performances (IOU, precision and recall) of predictions (N=5,094) between using the original training sets and image augmented training sets. Significant symbols are t-test results (ns: p > 0.05; *: p <= 0.05; **: p <= 0.01; ***: p <= 0.001; ****: p <= 0.0001).

**Figure S4.**
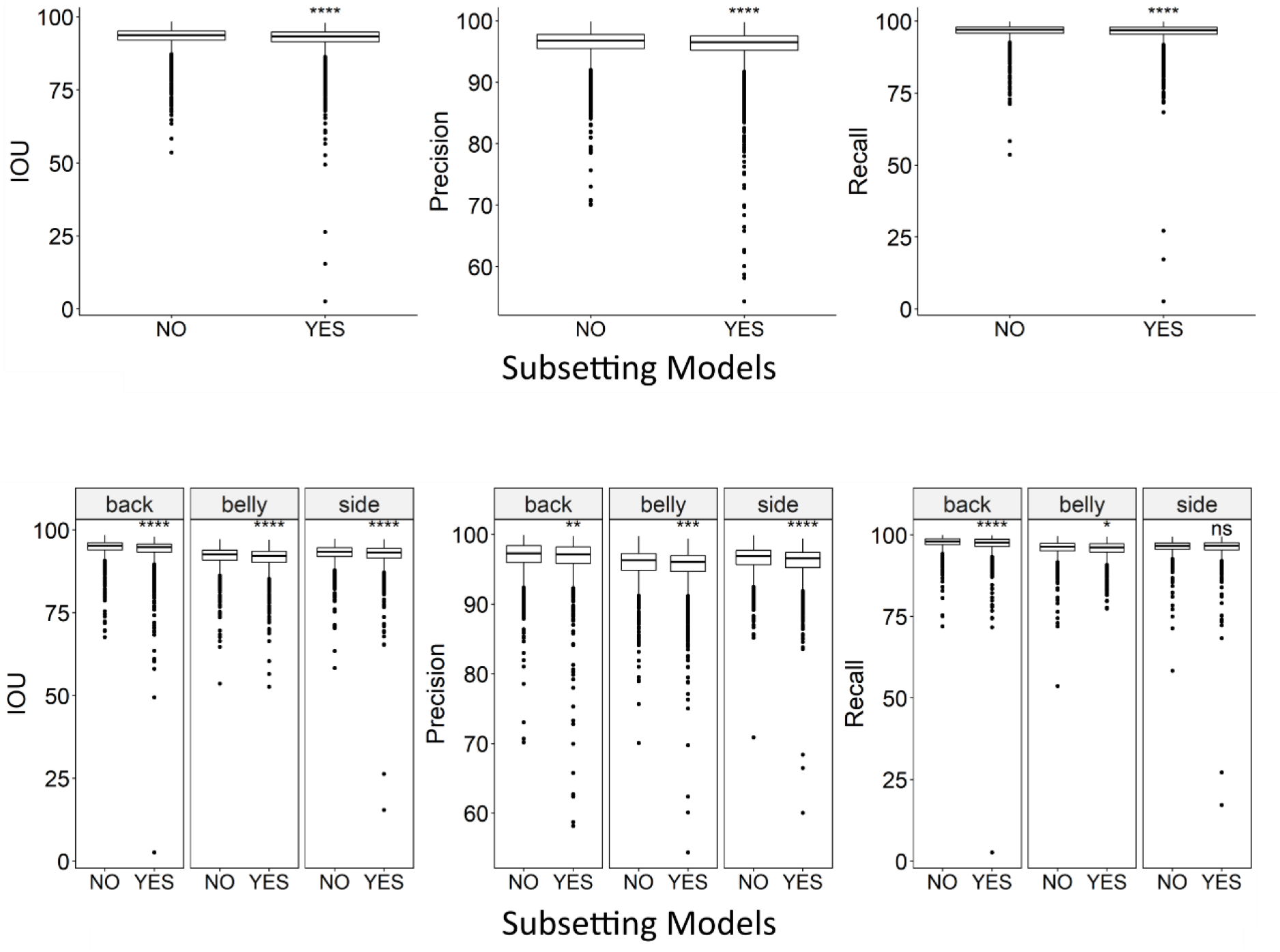
Boxplots of the performances (IOU, precision and recall) of predictions (N=5,094) between using one model and three separate models. Significant symbols are t-test results (ns: p > 0.05; *: p <= 0.05; **: p <= 0.01; ***: p <= 0.001; ****: p <= 0.0001).

**Figure S5.**
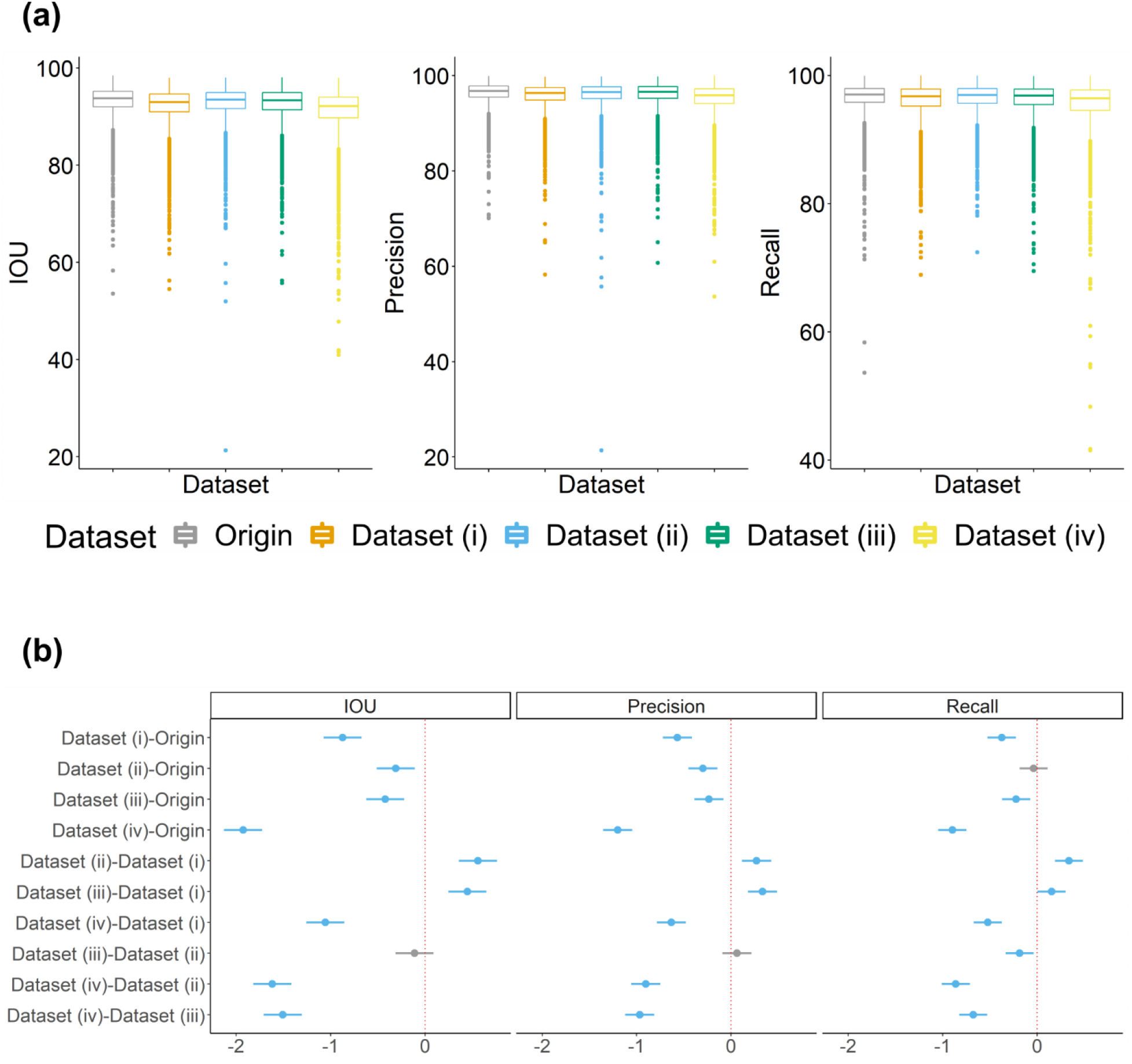
(a) Boxplots of the performances (IOU, precision and recall) of predictions (N=5,094) using the original dataset and low-quality datasets. Dataset (i) rotated (angles between −45 to 45), translated (−500 to 500 pixels on x and y axes) and scaled (scale ratio from 0.8 to 1.2) images; Dataset (ii) horizontal flip 50% images randomly; Dataset (iii) images with random contrast and brightness; Dataset (iv) the combination of (i), (ii) and (iii). (b) Plots of Tukey’s test (95% family-wise confidence level) on whether metric (IOU, precision and recall) differences among tested datasets are significantly different (blue: significance; grey: no significance) from 0 (red dotted lines).

**Figure S6.**
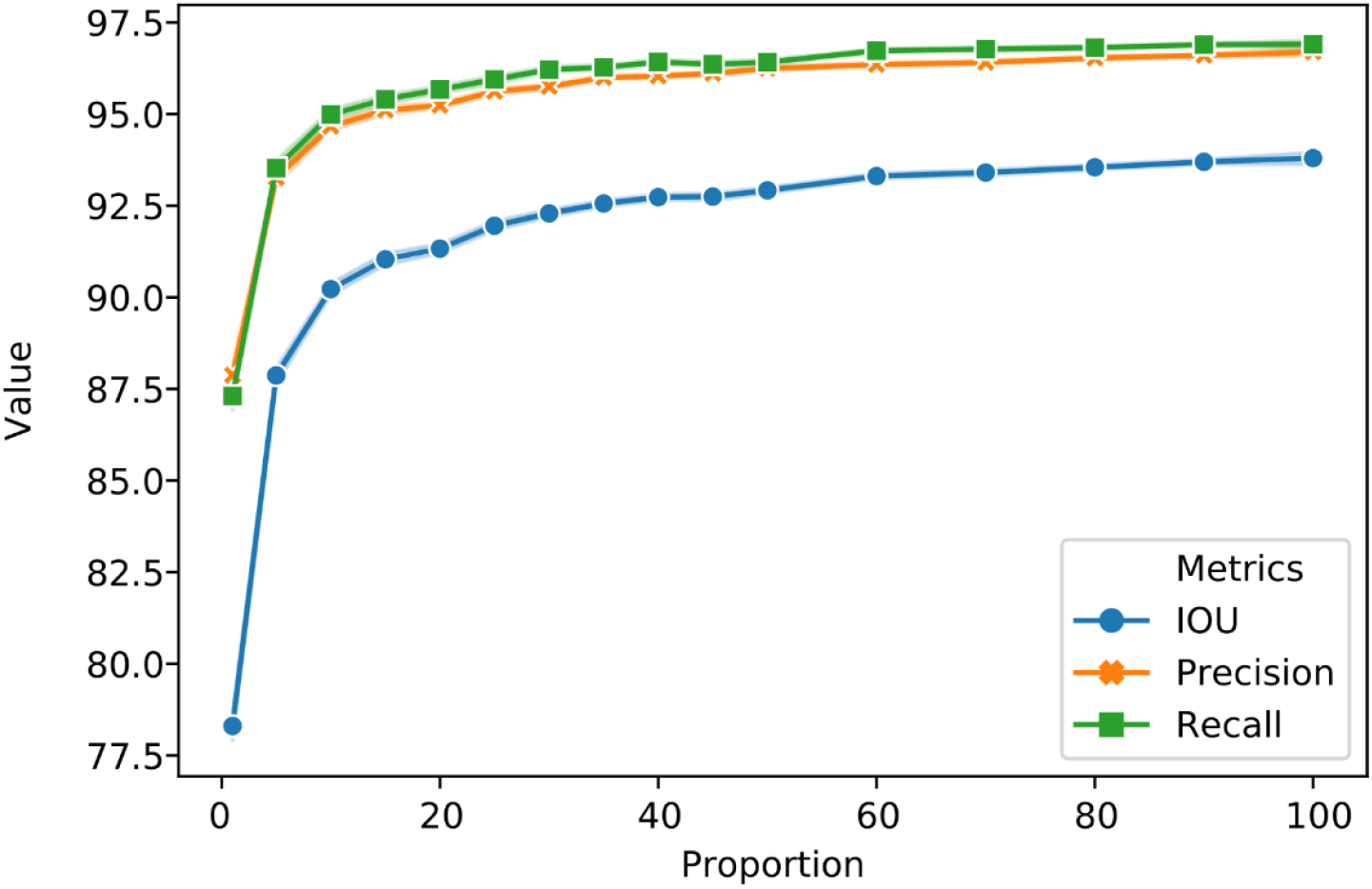
The performances (IOU, precision and recall) of the same validation set (N=1,018) using 15 proportions (1%, every 5% from 5% to 50% and every 10% from 50% to 90%) of the original training set.

**Figure S7.**
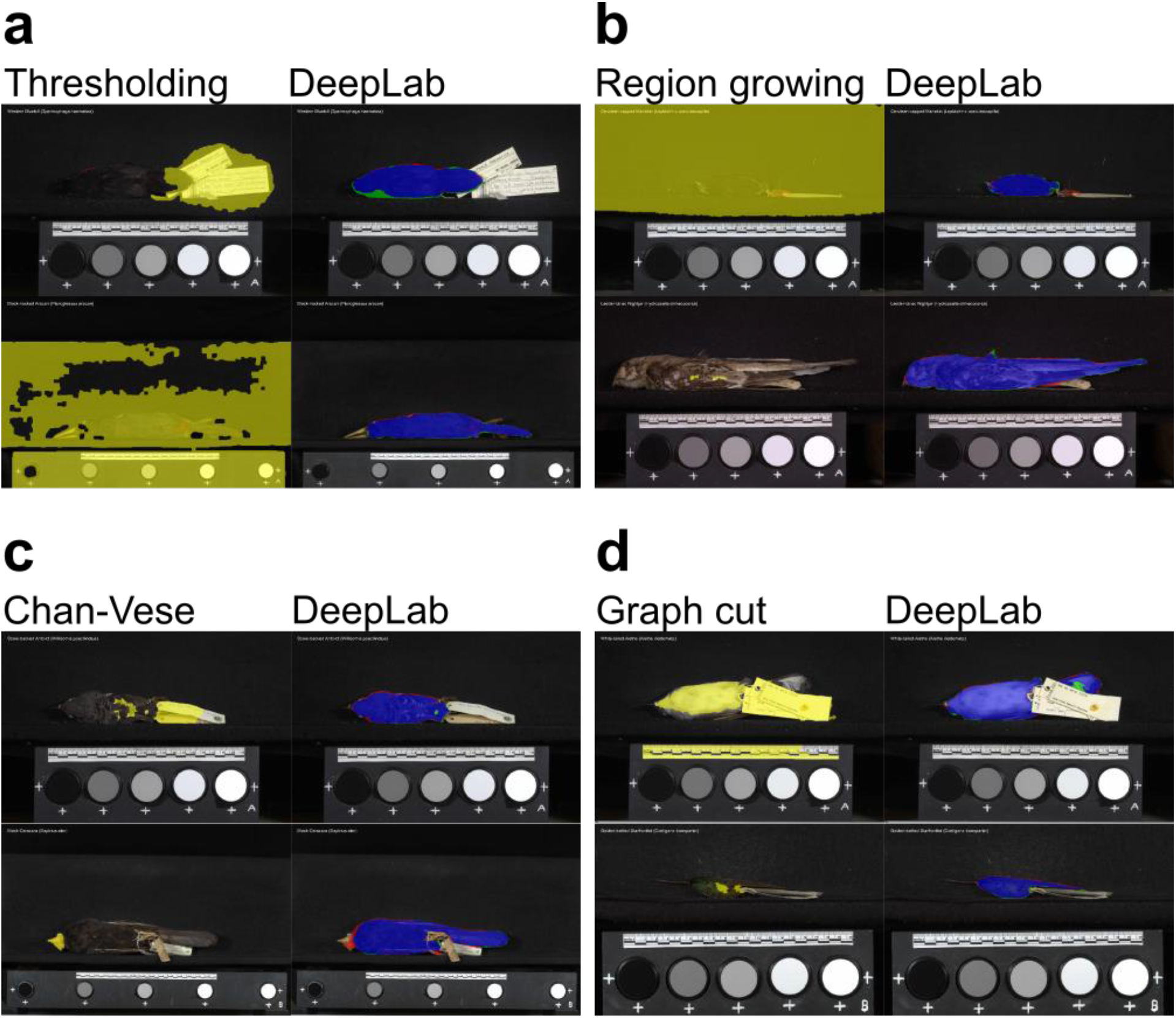
Examples of poorly segmented results from classic methods as well as deep learning predictions of the corresponding images. (a) Thresholding; (b) Region growing; (c) Chan-Vese; (d) Graph cut. Yellow is the segmentation from classic methods. Deep learning results are represented in blue, red and green as defined in Fig. 3.

**Figure S8.**
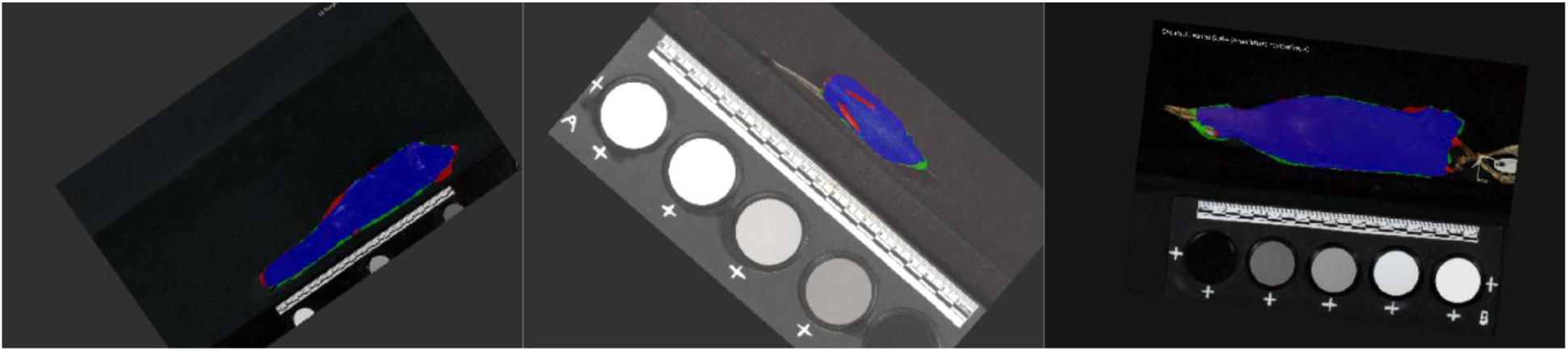
Examples of predictions using Dataset (iv) of the low-quality datasets.

## Supplementary Tables

**Table S1.** IOU, precision and recall of predictions from the DeepLabV3+ model

|  | MEAN | SD | MIN | MAX |
| --- | --- | --- | --- | --- |
| **IOU OVERALL (N=5094)** | 93.1 | 3.2 | 53.6 | 98.5 |
| **BACK (N=1698)** | 94.6 | 2.8 | 67.6 | 98.5 |
| **BELLY (N=1698)** | 91.9 | 3.4 | 53.6 | 97.2 |
| **SIDE (N=1698)** | 92.9 | 2.9 | 58.3 | 97.2 |
| **PRECISION OVERALL (N=5094)** | 96.3 | 2.4 | 70.1 | 99.9 |
| **BACK (N=1698)** | 96.8 | 2.5 | 70.2 | 99.9 |
| **BELLY (N=1698)** | 95.7 | 2.5 | 70.1 | 99.8 |
| **SIDE (N=1698)** | 96.4 | 2.0 | 70.9 | 99.9 |
| **RECALL OVERALL (N=5094)** | 96.6 | 2.5 | 53.6 | 99.9 |
| **BACK (N=1698)** | 97.6 | 2.0 | 72.0 | 99.9 |
| **BELLY (N=1698)** | 95.8 | 2.7 | 53.6 | 99.6 |
| **SIDE (N=1698)** | 96.2 | 2.4 | 58.3 | 99.4 |

**Table S2.**
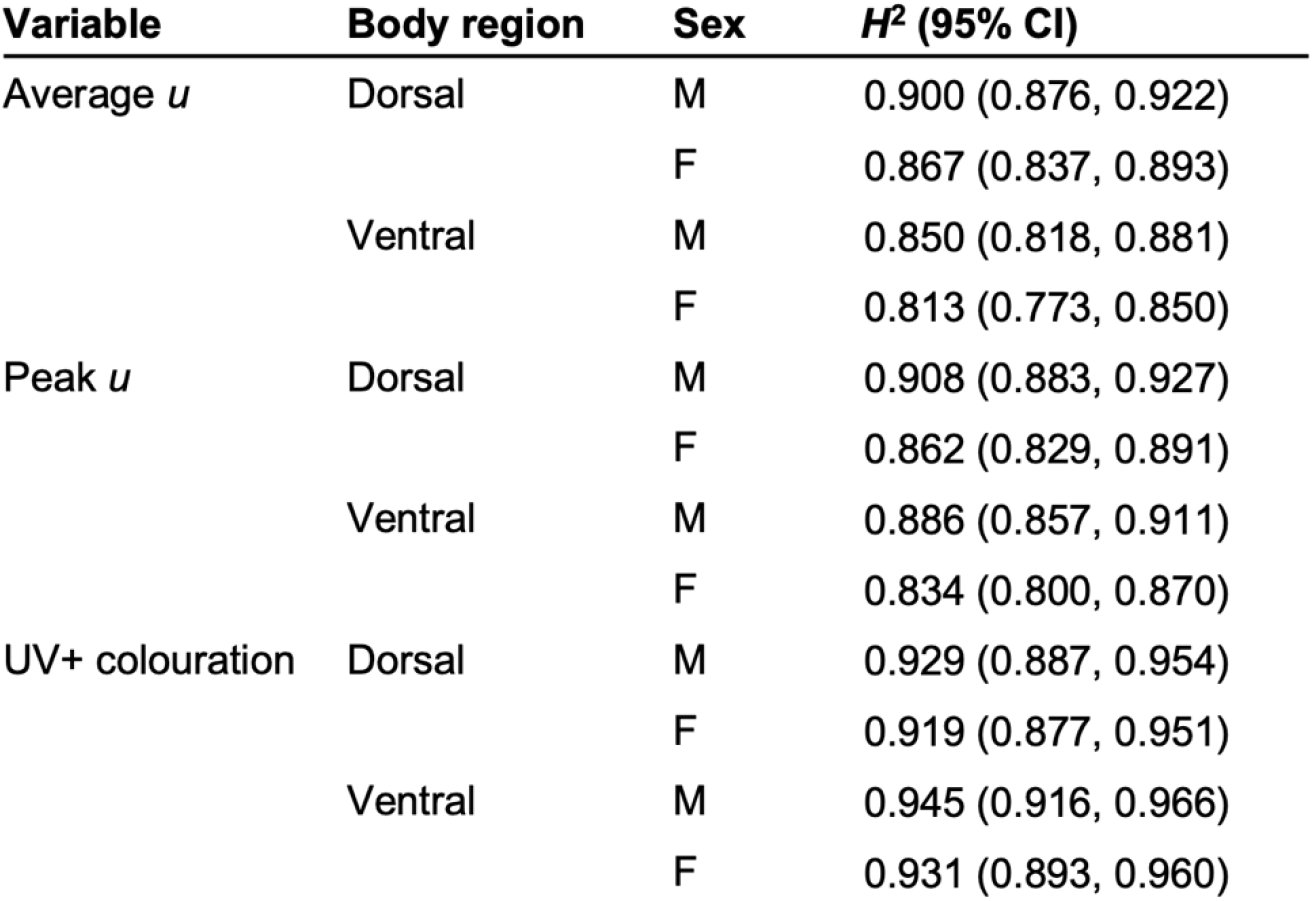
Phylogenetic heritability (H^2^) estimates for UV colouration metrics across passerine birds (n = 4,545). All models were run over 100 posterior phylogenetic trees.

| Variable | Body region | Sex | $H^2$ (95% CI) |
| --- | --- | --- | --- |
| Average $u$ | Dorsal | M | 0.900 (0.876, 0.922) |
|  |  | F | 0.867 (0.837, 0.893) |
|  | Ventral | M | 0.850 (0.818, 0.881) |
|  |  | F | 0.813 (0.773, 0.850) |
| Peak $u$ | Dorsal | M | 0.908 (0.883, 0.927) |
|  |  | F | 0.862 (0.829, 0.891) |
|  | Ventral | M | 0.886 (0.857, 0.911) |
|  |  | F | 0.834 (0.800, 0.870) |
| UV+ colouration | Dorsal | M | 0.929 (0.887, 0.954) |
|  |  | F | 0.919 (0.877, 0.951) |
|  | Ventral | M | 0.945 (0.916, 0.966) |
|  |  | F | 0.931 (0.893, 0.960) |

**Table S3.** Bayesian phylogenetic mixed model results for the effect of predictor variables on plumage UV reflectance in passerine species (n = 4,527). All variables were standardised (mean = 0, sd = 1) prior to model fitting. M, male; UVS, ultraviolet sensitive. *, P < 0.05; **, P < 0.01; ***, P < 0.001. All models were run over 100 posterior phylogenetic trees.

| Variable | Term | Estimate (95% CI) | $P_{\text{MCMC}}$ |
| --- | --- | --- | --- |
| Average $u$ | (Intercept) | -0.087 (-1.058, 0.877) | 0.858 |
|  | Body region<sub>[Ventral]</sub> | -0.259 (-0.267, -0.251) | <0.001*** |
|  | Sex<sub>[M]</sub> | 0.141 ( 0.132, 0.149) | <0.001*** |
|  | UVB | 0.046 ( 0.026, 0.065) | <0.001*** |
|  | Forest dependency<sub>[High]</sub> | 0.094 ( 0.072, 0.116) | <0.001*** |
|  | Visual system<sub>[UVS]</sub> | 0.150 (-0.117, 0.414) | 0.268 |
| Peak $u$ | (Intercept) | 0.055 (-0.900, 1.037) | 0.910 |
|  | Body region<sub>[Ventral]</sub> | -0.159 (-0.167, -0.152) | <0.001*** |
|  | Sex<sub>[M]</sub> | 0.175 ( 0.167, 0.183) | <0.001*** |
|  | UVB | 0.063 ( 0.044, 0.082) | <0.001*** |
|  | Forest dependency<sub>[High]</sub> | 0.114 ( 0.092, 0.136) | <0.001*** |
|  | Visual system<sub>[UVS]</sub> | 0.226 (-0.084, 0.537) | 0.153 |
| UV+ colouration | (Intercept) | -6.918 (-13.025, -0.943) | 0.023* |
|  | Body region<sub>[Ventral]</sub> | 0.431 ( 0.361, 0.501) | <0.001*** |
|  | Sex<sub>[M]</sub> | 0.464 ( 0.393, 0.533) | <0.001*** |
|  | UVB | 0.217 ( 0.047, 0.386) | 0.012* |
|  | Forest dependency<sub>[High]</sub> | 0.414 ( 0.238, 0.588) | <0.001*** |
|  | Visual system<sub>[UVS]</sub> | 0.886 ( -0.614, 2.420) | 0.247 |

## References

Abadi, M., A. Agarwal, P. Barham, E. Brevdo, Z. Chen, C. Citro, G. S. Corrado, A. Davis, J. Dean, M. Devin, S. Ghemawat, I. Goodfellow, A. Harp, G. Irving, M. Isard, Y. Jia, R. Jozefowicz, L. Kaiser, M. Kudlur, J. Levenberg, D. Mané, R. Monga, S. Moore, D. Murray, C. Olah, M. Schuster, J. Shlens, B. Steiner, I. Sutskever, K. Talwar, P. Tucker, V. Vanhoucke, V. Vasudevan, F. Viégas, O. Vinyals, P. Warden, M. Wattenberg, M. Wicke, Y. Yu, and Z. X. 2016. TensorFlow: large-scale machine learning on heterogeneous distributed systems. arXiv 1603:04467.

Adams, R. and L. Bischof. 1994. Seeded region growing. IEEE Transactions on Pattern Analysis and Machine Intelligence 18:641–647.

Aljabar, P., R. A. Heckemann, A. Hammers, J. V. Hajnal, and D. Rueckert. 2009. Multi-atlas based segmentation of brain images: atlas selection and its effect on accuracy. NeuroImage 46:726–738.

Andriluka, M., L. Pishchulin, P. Gehler, and B. Schiele. 2014. 2D human pose estimation: new benchmark and state of the art analysis. Pp. 3686–3693. 2014 IEEE Conference on Computer Vision and Pattern Recognition.

Baiker, M., J. Milles, J. Dijkstra, T. D. Henning, A. W. Weber, I. Que, E. L. Kaijzel, C. W. Löwik, J. H. Reiber, and B. P. Lelieveldt. 2010. Atlas-based whole-body segmentation of mice from low-contrast Micro-CT data. Med. Image Anal. 14:723–737.

Basu, S., S. Ganguly, S. Mukhopadhyay, R. DiBiano, M. Karki, and R. Nemani. 2015. DeepSat - a learning framework for satellite imagery. Pp. 1–10. Proceedings of the 23rd SIGSPATIAL International Conference on Advances in Geographic Information Systems.

Beckmann, M., T. Václavík, A. M. Manceur, L. Šprtová, H. von Wehrden, E. Welk, and A. F. Cord. 2014. glUV: a global UV-B radiation data set for macroecological studies. Methods in Ecology and Evolution 5:372–383.

Boykov, Y. Y. and M. P. Jolly. 2001. Interactive graph cuts for optimal boundary & region segmentation of objects in N-D images. Pp. 105–112. Proceedings Eighth IEEE International Conference on Computer Vision.

Bradski, G. 2000. The OpenCV Library. Dr. Dobb’s Journal of Software Tools 120:122–125.

Caro, T. and M. Koneru. 2021. Towards an ecology of protective coloration. Biological Reviews 96:611–641.

Chan, T. F. and L. A. Vese. 2001. Active contours without edges. IEEE Transactions on Image Processing 10:266–277.

Chang, Y. L. and X. Li. 1994. Adaptive image region-growing. IEEE Transactions on Image Processing 3:868 – 872.

Chen, L. C., G. Papandreou, I. Kokkinos, K. Murphy, and A. L. Yuille. 2017a. DeepLab: semantic image segmentation with deep convolutional nets, atrous convolution, and fully connected CRFs. arXiv 1606:00915.

Chen, L. C., G. Papandreou, F. Schroff, and H. Adam. 2017b. Rethinking atrous convolution for semantic image segmentation. arXiv 1706:05587.

Chen, L. C., Y. Zhu, G. Papandreou, F. Schroff, and H. Adam. 2018. Encoder-decoder with atrous separable convolution for semantic image segmentation. arXiv 1802:02611.

Christin, S., É. Hervet, and N. Lecomte. 2019. Applications for deep learning in ecology. Methods in Ecology and Evolution 10:1632–1644.

Coffin, D. 2016. DCRAW V. 9.27 https://www.cybercom.net/~dcoffin/dcraw/.

Cooney, C. R., J. A. Bright, E. J. R. Capp, A. M. Chira, E. C. Hughes, C. J. A. Moody, L. O. Nouri, Z. K. Varley, and G. H. Thomas. 2017. Mega-evolutionary dynamics of the adaptive radiation of birds. Nature 542:344–347.

Cooney, C. R., Z. K. Varley, L. O. Nouri, C. J. A. Moody, M. D. Jardine, and G. H. Thomas. 2019. Sexual selection predicts the rate and direction of colour divergence in a large avian radiation. Nature communications 10:1773.

Cordts, M., M. Omran, S. Ramos, R. T., M. Enzweiler, R. Benenson, U. Franke, S. Roth, and B. Schiele. 2016. The Cityscapes dataset for semantic urban scene understanding. arXiv 1604:01685.

Cuthill, I. C., W. L. Allen, K. Arbuckle, B. Caspers, G. Chaplin, M. E. Hauber, G. E. Hill, N. G. Jablonski, C. D. Jiggins, A. Kelber, J. Mappes, J. Marshall, R. Merrill, D. Osorio, R. Prum, N. W. Roberts, A. Roulin, H. M. Rowland, T. N. Sherratt, J. Skelhorn, M. P. Speed, M. Stevens, M. C. Stoddard, D. Stuart-Fox, L. Talas, E. Tibbetts, and T. Caro. 2017. The biology of color. Science 357:eaan0221.

Dale, J., C. J. Dey, K. Delhey, B. Kempenaers, and M. Valcu. 2015. The effects of life history and sexual selection on male and female plumage colouration. Nature 527:367–370.

Delhey, K. 2020. Revealing the colourful side of birds: spatial distribution of conspicuous plumage colours on the body of Australian birds. J. Avian Biol. 51:e02222.

Deng, J., W. Dong, R. Socher, L. J. Li, K. Li, and L. Fei-Fei. 2009. ImageNet: a large-scale hierarchical image database. 2009 IEEE Conference on Computer Vision and Pattern Recognition.

Endler, J. A. 1992. Signals, signal conditions, and the direction of evolution. Am. Nat. 139:S125–S153.

Endler, J. A. 1993a. The color of light in forests and its implications. Ecol. Monogr. 63:1–27.

Endler, J. A. 1993b. Some general comments on the evolution and design of animal communication systems. Philos. Trans. R. Soc. London Ser. B 340:215–225.

Everingham, M., S. M. A. Eslami, L. Van Gool, C. K. I. Williams, J. Winn, and A. Zisserman. 2015. The PASCAL Visual Object Classes challenge – a retrospective. International Journal of Computer Vision 111:98–136.

Fan, J., D. K. Y. Yau, A. K. Elmagarmid, and W. G. Aref. 2001. Automatic image segmentation by integrating color-edge extraction and seeded region growing. IEEE Transactions on Image Processing 10:1454–1466.

Felice, R. N. and A. Goswami. 2018. Developmental origins of mosaic evolution in the avian cranium. Proc. Natl. Acad. Sci. U.S.A. 15:555–560.

Gomez, D. and M. Théry. 2007. Simultaneous crypsis and conspicuousness in color patterns: comparative analysis of a Neotropical rainforest bird community. Am. Nat. 169:S42–S61.

Hadfield, J. D. 2010. MCMC methods for multi-response generalised linear mixed models: the MCMCglmm R package. Journal of Statistical Software 33:1–22.

Hadfield, J. D. and S. Nakagawa. 2010. General quantitative genetic methods for comparative biology: phylogenies, taxonomies and multi-trait models for continuous and categorical characters. J. Evol. Biol. 23:494–508.

Haralick, R. M., S. R. Sternberg, and X. Zhuang. 1987. Image analysis using mathematical morphology. IEEE Transactions on Pattern Analysis and Machine Intelligence 9:532–550.

Håstad, O., J. Victorsson, and A. Ödeen. 2005. Differences in color vision make passerines less conspicuous in the eyes of their predators. Proc. Natl. Acad. Sci. U.S.A. 102:6391–6394.

Hausmann, F., K. E. Arnold, N. J. Marshall, and I. P. Owens. 2003. Ultraviolet signals in birds are special. Proc. R. Soc. London Ser. B 270:61–67.

He, K., X. Zhang, S. Ren, and J. Sun. 2016. Deep residual learning for image recognition. Pp. 770–778. 2016 IEEE Conference on Computer Vision and Pattern Recognition.

Healy, K., T. Guillerme, S. Finlay, A. Kane, S. B. Kelly, D. McClean, D. J. Kelly, I. Donohue, A. L. Jackson, and N. Cooper. 2014. Ecology and mode-of-life explain lifespan variation in birds and mammals. Proc. R. Soc. London Ser. B 281:20140298.

Hestness, J., S. Narang, N. Ardalani, G. Diamos, H. Jun, H. Kianinejad, M. M. A. Patwary, Y. Yang, and Y. Zhou. 2017. Deep learning scaling is predictable, empirically. arXiv 1712:00409.

Hijmans, R. J. 2020. raster: geographic data analysis and modeling. R package version 3.4-5. https://CRAN.R-project.org/package=raster.

Hinton, G. E., N. Srivastava, A. Krizhevsky, I. Sutskever, and R. R. Salakhutdinov. 2012. Improving neural networks by preventing co-adaptation of feature detectors. arXiv 1207:0580.

Hudson, L. N., V. Blagoderov, A. Heaton, P. Holtzhausen, L. Livermore, B. W. Price, S. van der Walt, and V. S. Smith. 2015. Inselect: automating the digitization of natural history collections. PloS one 10:e0143402.

Hussein, B. R., O. A. Malik, W.-H. Ong, and J. W. F. Slik. 2020. Semantic segmentation of herbarium specimens using deep learning techniques. Pp. 321–330. Computational Science and Technology.

Jetz, W., G. H. Thomas, J. B. Joy, K. Hartmann, and A. O. Mooers. 2012. The global diversity of birds in space and time. Nature 491:444–448.

Joulin, A., L. van der Maaten, A. Jabri, and N. Vasilache. 2015. Learning visual features from large weakly supervised data. arXiv 1511:02251.

Kass, M., A. Witkin, and D. Terzopoulos. 1988. Snakes: active contour models. International Journal of Computer Vision 1:321–331.

Kingma, D. P. and J. L. Ba. 2014. ADAM: a method for stochastic optimisation. arXiv 1412:6980.

Kohler, R. 1981. A segmentation system based on thresholding. Computer Graphics and Image Processing 15:319–338.

Krizhevsky, A., I. Sutskever, and G. E. Hinton. 2012. ImageNet classification with deep convolutional neural networks. Pp. 1–9. Advances In Neural Information Processing Systems.

Kumar, Y. H. S., N. Manohar, and H. K. Chethan. 2015. Animal classification system: a block based approach. Procedia Computer Science 45:336–343.

Lee, J. S. 1983. Digital image smoothing and the signam filter. Computer Vision, Graphics, and Image Processing 24:255–269.

Lind, O. and K. Delhey. 2015. Visual modelling suggests a weak relationship between the evolution of ultraviolet vision and plumage coloration in birds. J. Evol. Biol. 28:715–722.

Lind, O., M. J. Henze, A. Kelber, and D. Osorio. 2017. Coevolution of coloration and colour vision? Philos. Trans. R. Soc. London Ser. B 372:20160338.

Long, J., E. Shelhamer, and T. Darrell. 2015. Fully convolutional networks for semantic segmentation. Pp. 3431–3440. 2016 IEEE Conference on Computer Vision and Pattern Recognition.

Loshchilov, I. and F. Hutter. 2016. SGDR: stochastic gradient descent with warm restarts. arXiv 1608:03983.

Lürig, M. D., S. Donoughe, E. I. Svensson, A. Porto, and M. Tsuboi. 2021. Computer vision, machine learning, and the promise of phenomics in ecology and evolutionary biology. Frontiers in Ecology and Evolution 9:642774.

Lynch, M. 1991. Methods for the analysis of comparative data in evolutionary biology. Evolution 45:1065–1080.

Maia, R., H. Gruson, J. A. Endler, T. E. White, and R. B. O’Hara. 2019. pavo 2: new tools for the spectral and spatial analysis of colour in R. Methods in Ecology and Evolution 10:1097–1107.

Maia, R., D. R. Rubenstein, and M. D. Shawkey. 2013. Key ornamental innovations facilitate diversification in an avian radiation. Proc. Natl. Acad. Sci. U.S.A. 110:10687–10692.

Meijering, E. 2012. Cell segmentation: 50 years down the road. IEEE Signal Processing Magazine 29:140–145.

Milioto, A., P. Lottes, and C. Stachniss. 2018. Real-time semantic segmentation of crop and weed for precision agriculture robots leveraging background knowledge in CNNs. Pp. 2229–2235. 2018 IEEE International Conference on Robotics and Automation.

Miller, E. T., G. M. Leighton, B. G. Freeman, A. C. Lees, and R. A. Ligon. 2019. Ecological and geographical overlap drive plumage evolution and mimicry in woodpeckers. Nature communications 10:1602.

Newell, A., K. Yang, and J. Deng. 2016. Stacked hourglass networks for human pose estimation. arXiv 1603:06937.

Nicolaï, M. P. J., M. D. Shawkey, S. Porchetta, R. Claus, and L. D’Alba. 2020. Exposure to UV radiance predicts repeated evolution of concealed black skin in birds. Nature communications 11:2414.

Ödeen, A. and O. Håstad. 2013. The phylogenetic distribution of ultraviolet sensitivity in birds. BMC Evol. Biol. 13:36.

Ödeen, A., O. Håstad, and P. Alström. 2011. Evolution of ultraviolet vision in the largest avian radiation - the passerines. BMC Evol. Biol. 11:313.

Ödeen, A., S. Pruett-Jones, A. C. Driskell, J. K. Armenta, and O. Hastad. 2012. Multiple shifts between violet and ultraviolet vision in a family of passerine birds with associated changes in plumage coloration. Proc. R. Soc. London Ser. B 279:1269–1276.

Otsu, N. 1979. A threshold selection method from gray-level histograms. IEEE Transactions on Systems, Man, and Cybernetics 9:62–66.

Potena, C., D. Nardi, and A. Pretto. 2017. Fast and accurate crop and weed identification with summarized train sets for precision agriculture in W. Chen, K. Hosoda, E. Menegatti, M. Shimizu, and H. Wang, eds. Intelligent Autonomous Systems 14. IAS 2016. Advances in Intelligent Systems and Computing, vol 531. Springer, Cham.

Ruder, S. 2016. An overview of gradient descent optimization algorithms. arXiv 1609:04747.

Schliep, K. P. 2011. phangorn: phylogenetic analysis in R. Bioinformatics 27:592–593.

Sezgin, M. and B. Sankur. 2004. Survey over image thresholding techniques and quantitative performance evaluation. Journal of Electronic Imaging 13:146–165.

Sheard, C., M. H. C. Neate-Clegg, N. Alioravainen, S. E. I. Jones, C. Vincent, H. E. A. MacGregor, T. P. Bregman, S. Claramunt, and J. A. Tobias. 2020. Ecological drivers of global gradients in avian dispersal inferred from wing morphology. Nature communications 11:2463.

Stevens, M. and I. C. Cuthill. 2007. Hidden messages: are ultraviolet signals a special channel in avian communication? Bioscience 57:501–507.

Stoddard, M. C. and R. O. Prum. 2008. Evolution of avian plumage color in a tetrahedral color space: a phylogenetic analysis of New World buntings. Am. Nat. 171:755–776.

Stoddard, M. C. and R. O. Prum. 2011. How colorful are birds? Evolution of the avian plumage color gamut. Behav. Ecol. 22:1042–1052.

Szegedy, C., W. Liu, Y. Jia, P. Sermanet, S. Reed, D. Anguelov, D. Erhan, V. Vanhoucke, and A. Rabinovich. 2014. Going deeper with convolutions. arXiv 1409:4842.

Troscianko, J. and M. Stevens. 2015. Image calibration and analysis toolbox – a free software suite for objectively measuring reflectance, colour and pattern. Methods in Ecology and Evolution 6:1320–1331.

Unger, J., D. Merhof, and S. Renner. 2016. Computer vision applied to herbarium specimens of German trees: testing the future utility of the millions of herbarium specimen images for automated identification. BMC Evol. Biol. 16:248.

Wei, S. E., V. Ramakrishna, T. Kanade, and Y. Sheikh. 2016. Convolutional pose machines. Pp. 4724–4732. 2016 IEEE Conference on Computer Vision and Pattern Recognition.

